# SMARCA4 MUTATIONS PROMOTE CHEMOTHERAPY RESISTANCE IN T-ACUTE LYMPHOBLASTIC LEUKEMIA THROUGH ALTERED CHROMATIN REMODELING AT AP-1-ASSOCIATED REGULATORY REGIONS

**DOI:** 10.64898/2026.11.01.763277

**Authors:** Violeta García-Hernández, Maria Maqueda, Giulia Faccincani, Alba Vidal, Jessica González, Arnau Iglesias, Marina Alcaide-Miranda, Cinzia Benetti, Eleonora Ponzo, Francesco Boccalatte, Sarah J Bray, Lluis Espinosa, Anna Bigas

**Author notes:** Corresponding authors:* Anna Bigas and Violeta García-Hernández Cancer Research Program. CIBERONC. Hospital del Mar Research Institute Barcelona Doctor Aiguader 88, 08003, Barcelona, Spain.

## Abstract

Chromatin remodelers play a critical role in orchestrating chemotherapy responses in cancer cells. Components of the SWI/SNF complex are frequently mutated across cancer types, with the ATPase subunit SMARCA4 among the most commonly altered. In T-cell acute lymphoblastic leukemia (T-ALL), however, SMARCA4 mutations are rare at diagnosis but enriched in relapse samples, suggesting a potential role in therapy resistance. Using isogenic T-ALL cell lines with or without SMARCA4 mutations, we show that mutant SMARCA4 confers resistance to a broad range of chemotherapeutic agents. By combining ATAC-seq and RNA- seq, we demonstrate that SMARCA4-mutant cells display an attenuated transcriptional response to chemotherapy accompanied by impaired remodeling of AP-1-associated regulatory regions linked to stress response. These cells downregulate stress-related genes while maintaining expression programs associated with cellular fitness. Pharmacological inhibition of AP-1 phenocopies this response, further supporting a functional link between SWI/SNF-mediated remodeling at AP-1-associated regulatory regions, AP-1 activity, and chemotherapy sensitivity. Together, our findings indicate that SWI/SNF-mediated remodeling of these regions contributes to an effective chemotherapy-induced stress response, and that its failure provides T-ALL cells with a survival advantage that contributes to relapse-associated therapy resistance.

## INTRODUCTION

T-cell Acute Lymphoblastic Leukemia (T-ALL) is an aggressive hematological malignancy characterized by the outgrowth of T lymphoid progenitors. Although it is a relatively rare disease (0.6 in 100,000 per year) affecting both children and adults and treatment outcomes have significantly improved in pediatric patients^1,2^, relapse remains a major clinical challenge, occurring in approximately 15% of children and 50% of adult patients^1,3,4^. Importantly, patients with relapse T-ALL have very limited therapeutic options and poor prognosis. Understanding how leukemic cells acquire resistance to therapy is therefore a central research goal for improving survival in leukemia and other cancers. While genetic alterations and cellular context clearly influence the response to chemotherapy, increasing evidence indicates that cellular plasticity driven by epigenetic mechanisms plays a key role in disease persistence and relapse^5–7^.

Chromatin remodeling is a fundamental regulatory mechanism that controls access to the genome by dynamically altering nucleosome positioning and chromatin structure^8^. Among the major chromatin remodeling complexes, the SWI/SNF (SWItch/Sucrose Non-Fermentable) family plays a crucial role in modulating transcription, DNA repair, replication, and cell differentiation through ATP-dependent remodeling of nucleosomes^9,10^. Central to the function of many SWI/SNF complexes is the catalytic ATPase subunit SMARCA4 (also known as BRG1), which provides the energy required to reposition or evict nucleosomes, thereby influencing gene accessibility and transcriptional regulation. SMARCA4-mediated chromatin remodeling is essential for normal cellular processes^11–13^, and its dysregulation has been implicated in numerous diseases, particularly cancer, where mutations or loss of function can disrupt gene expression programs and cellular identity^10,13–16^. Understanding the molecular mechanisms by which SWI/SNF complexes and SMARCA4 regulate chromatin architecture is therefore critical for elucidating how epigenetic control contributes to development and disease.

In T-ALL, mutations affecting chromatin remodeling factors are not among the most frequent alterations^17^; however, SMARCA4 mutations are detected in approximately 10% of relapse patient samples^18^. Notably, these alterations are typically heterozygous and affect the ATPase domain, suggesting impaired enzymatic activity^14,19^ or, as previously proposed^20^, a potential dominant negative function that interferes with normal SWI/SNF complex activity. This observation is particularly intriguing because while SMARCA4 can function as a leukemogenic factor in acute myeloid leukemia (AML)^13,21,22^ and has also been implicated in T-ALL maintenance^23^, the presence of hypomorphic mutations in relapse T-ALL suggests that they may confer a selective advantage during disease progression or therapy exposure.

These observations raise important questions regarding the role of SMARCA4-mediated chromatin remodeling in leukemic cell plasticity and therapeutic resistance. Despite growing interest in SWI/SNF function in leukemia^13,21–23^, the consequences of SMARCA4 mutations in hematological malignancies are poorly defined, and the few studies addressing SWI/SNF impairment in the context of therapy exposure have relied on depletion models^24,25^. In particular, it remains unclear how alterations in SMARCA4 function reshape chromatin accessibility and transcriptional programs in T-ALL cells, and whether these changes promote adaptive states that allow leukemic cells to survive chemotherapy. Elucidating the mechanisms by which SMARCA4 mutations contribute to relapse may therefore reveal critical epigenetic vulnerabilities and identify new strategies to overcome treatment resistance in T-ALL.

We have now dissected the effect of SMARCA4 mutations in T-ALL by engineering isogenic mutant and wildtype cell lines and demonstrated that SMARCA4 mutations confer resistance to different chemotherapy agents in vitro and in vivo. We show how chromatin remodeling mechanisms are at the base of this resistance affecting the accessibility at AP-1-dependent regulatory elements that control stress responses. This defect limits chemotherapy-induced transcriptional reprogram ming resulting in attenuated activation of stress-response genes with relative preservation of fitness-associated gene expression.

## RESULTS

### Heterozygous SMARCA4 mutations targeting the ATPase domain are enriched in relapsed T-ALL

By analyzing relapse mutations in five pediatric and adult T-ALL cohorts (n = 97)^18,26–29^, we now found that approximately 10% of relapsed cases carry SMARCA4 mutations (Table S1). These alterations were heterozygous and clustered within the ATPase domain of SMARCA4, including both the N- and C-terminal regions (Figure 1A). Some of these mutations are predicted to have reduced enzymatic activity and have shown a dominant-negative effect on SWI/SNF complex function^14,19,20^.

**Figure 1.**
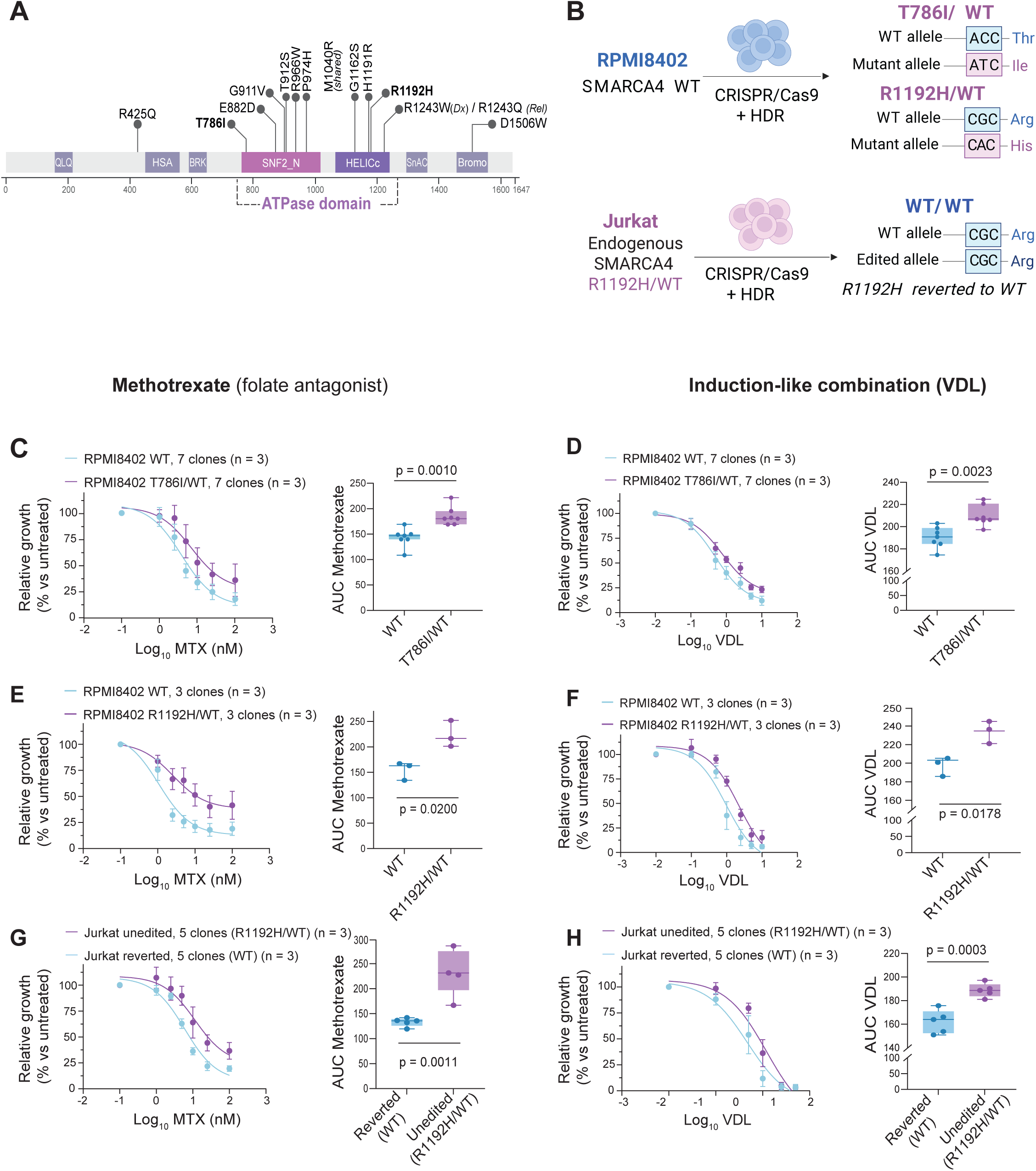
Relapse-associated mutations in the chromatin remodeler SMARCA4 promote chemotherapy resistance in T-ALL cell lines. **A.** SMARCA4 protein structure (UniProt, SMCA4_HUMAN) with the distribution of the mutations found in relapsed T-ALL cohorts. Mutations selected for gene editing in T-ALL cell lines are highlighted. QLQ: QLQ (Gln, Leu, Gln) motif; HSA: helicase/SANT-associated domain; BRK: brahma and kismet domain; SNF2-N: SNF2 family N-terminal domain, and HELICc: helicase superfamily C-terminal domain; SnAC: Snf2-ATP coupling, chromatin- remodeling complex domain; Bromo: bromodomain. **B.** Schematic overview of the generation of isogenic T-ALL cell lines carrying relapse- associated SMARCA4 mutations. Heterozygous T786I and R1192H mutations were introduced into RPMI8402 cells by CRISPR/Cas9-mediated homology-directed repair (HDR). The endogenous R1192H mutation present in Jurkat cells was reverted to the WT sequence. **C-D.** Sensitivity of RPMI8402 WT and T786I/WT clones to methotrexate (MTX; **C**) and induction-like chemotherapy (VDL; vincristine, dexamethasone and L-asparaginase; **D**). Left, dose-response curves. Right, area under the curve (AUC) quantification. **E-F.** Sensitivity of RPMI8402 WT and R1192H/WT clones to MTX (**E**) and VDL (**F**). **G–H.** Sensitivity of Jurkat unedited (R1192H/WT) and reverted WT clones to MTX (**G**) and VDL (**H**). For panels C–H, dose–response curves represent the mean ± SD of the indicated number of independent isogenic clones per genotype. For each clone, values at each drug concentration were calculated as the mean of three independent experiments. In the AUC plots, each dot represents an independent isogenic clone, with AUC values calculated as the mean of three independent experiments. Box plots show the minimum to maximum range, with the central line indicating the median and the box representing the 25th–75th percentiles. Statistical significance was assessed using two-tailed unpaired t-tests.

In contrast, SMARCA4 mutations were rare at diagnosis. SMARCA4 missense variants affected only 1.15% of cases in an independent diagnostic cohort (n = 1,309 T-ALL cases with genomic data at diagnosis^17^), and only one patient within the relapse cohorts carried detectable SMARCA4 mutations at diagnosis (Table S1). Relapse-associated SMARCA4 mutations are generally not detected in matched diagnostic samples. However, by mathematical modeling of clonal sizes in our adult cohort, we previously inferred that the relapse mutations were present at diagnosis in most of the patients (16 of 19 cases), albeit at frequencies below 0.01% and therefore beyond the detection limits of conventional sequencing approaches^18^. Supporting this notion, in the Kunz et al. cohort, one relapse-specific SMARCA4 mutation was only detected at diagnosis using highly sensitive targeted ultra-deep sequencing, at 0.83% Variant Allele Frequency (VAF)^26^. These observations are consistent with expansion or selection of SMARCA4-mutant subclones during treatment and relapse evolution.

### Relapse-associated mutations in the chromatin remodeler SMARCA4 promote chemotherapy resistance in T-ALL cell lines

To investigate the functional impact of relapse-associated mutations, we engineered human T- ALL cell lines with clinically observed SMARCA4 mutations using CRISPR/Cas9-mediated genome editing by an optimized homologous recombination strategy (Figure 1B).

We focused on two representative mutations affecting distinct conserved regions within the ATPase domain: T786I, located within the Walker A motif involved in ATP binding^20^, and R1192H, which has been linked to ATP–DNA coupling^14^. We introduced these mutations in heterozygosity into RPMI8402 cells, a cortical-like TAL1/LMO1-driven T-ALL model carrying the SIL-TAL1 translocation lacking alterations in SWI/SNF complex components (DepMap v26Q3; depmap.org^30^) and expressing high levels of SMARCA4 (Figure S1A). In parallel, we reverted the endogenous heterozygous R1192H mutation present in Jurkat cells (a relapse- derived TAL1-associated T-ALL cell line with a more mature phenotype) to the wild-type (WT) allele (Figure 1B). Although Jurkat cells also carry an additional heterozygous ATPase-domain variant (R1244H), this alteration is predicted to be benign (PolyPhen score = 0.035) and was not further manipulated. For each condition, we isolated isogenic single-cell–derived WT and mutant clones and confirmed SMARCA4 mutational status by Sanger sequencing of genomic DNA and mRNA transcripts (Figure S1B). SMARCA4 expression remained unchanged following introduction or reversion of heterozygous mutations, without compensatory upregulation of the alternative SWI/SNF ATPase SMARCA2, which is expressed at low levels in these cell lines (Figure S1C).

We next assessed the effects of SMARCA4 mutations on cell proliferation and viability, as acute inhibition or degradation of SMARCA4 has been shown to impair cell growth and survival in T-ALL models^23^. Despite some clonal variability, heterozygous SMARCA4-mutant cell lines did not exhibit major alterations in proliferation, cell-cycle distribution, or viability under basal culture conditions compared with WT controls (Figures S1D, S1F, S1H). In contrast, SMARCA4 knockdown in RPMI8402 cells profoundly reduced proliferation and viability (Figure S1E, S1G), in line with previous reports of acute SMARCA4 depletion^22,23^. These findings indicate that relapse-associated ATPase-domain mutations do not fully phenocopy SMARCA4 loss, and highlight a context-dependent role for SMARCA4 in leukemia maintenance.

We then evaluated the impact of SMARCA4 mutations on treatment response, using standard chemotherapy for T-ALL management. RPMI8402 clones carrying heterozygous SMARCA4 mutations (T786I/WT and R1192H/WT) consistently showed reduced sensitivity to multiple chemotherapy treatments, including methotrexate (MTX) and an induction-like chemotherapy combination (VDL: vincristine, dexamethasone, and L-asparaginase) (Figure 1C-1F), whereas the reversion of the R1192H mutation in Jurkat sensitized the cells to these treatments (Figure 1G-1H). We similarly found that RPMI8402 mutant cells were more resistant to the treatment with additional chemotherapeutic drugs, including etoposide and doxorubicin (Figure S1I).

SMARCA4-knockdown RPMI8402 cells also showed reduced response to different chemotherapy treatments (MTX, VDL, etoposide) (Figure S1J-K), although in this case, this effect may be partially influenced by the marked proliferation defect induced by acute SMARCA4 depletion.

### Relapse-associated SMARCA4 mutations confer a competitive advantage under chemotherapy pressure

We previously estimated that relapse-associated mutations often pre-exist at diagnosis in minor subclones^18^, supporting a model where therapy-sensitive and resistant populations coexist within the primary tumor and compete during treatment.

To model clonal competition during therapy, we mixed fluorescently labeled WT or SMARCA4- mutant cells at a 1:1 ratio and tracked population dynamics during chemotherapy exposure (Figure 2A). Under MTX or induction-like VDL treatment, RPMI8402 SMARCA4-mutant (T786I/WT) cells progressively outcompeted their WT counterparts in vitro (Figure 2B, 2C). This phenotype was reproducible across independent mutant clones and fluorescent labeling strategies (Figure S2A-S2B), and was similarly observed in the Jurkat model (Figure S2D, S2E). Importantly, the competitive advantage of SMARCA4-mutant cells emerged specifically under therapeutic pressure, and was not observed in untreated cultures, where mutant and WT cells showed comparable proliferative capacity, or even reduced in the case of Jurkat models (Figure 2D, S2E). This context-dependent phenotype indicates that the mutation does not simply increase baseline proliferation but rather enhances tolerance to therapeutic stress. Consistent with this interpretation, SMARCA4-mutant cells underwent reduced apoptosis following chemotherapy exposure, as measured by Annexin V staining after 72 h of MTX or VDL treatment (Figure 2E).

**Figure 2.**
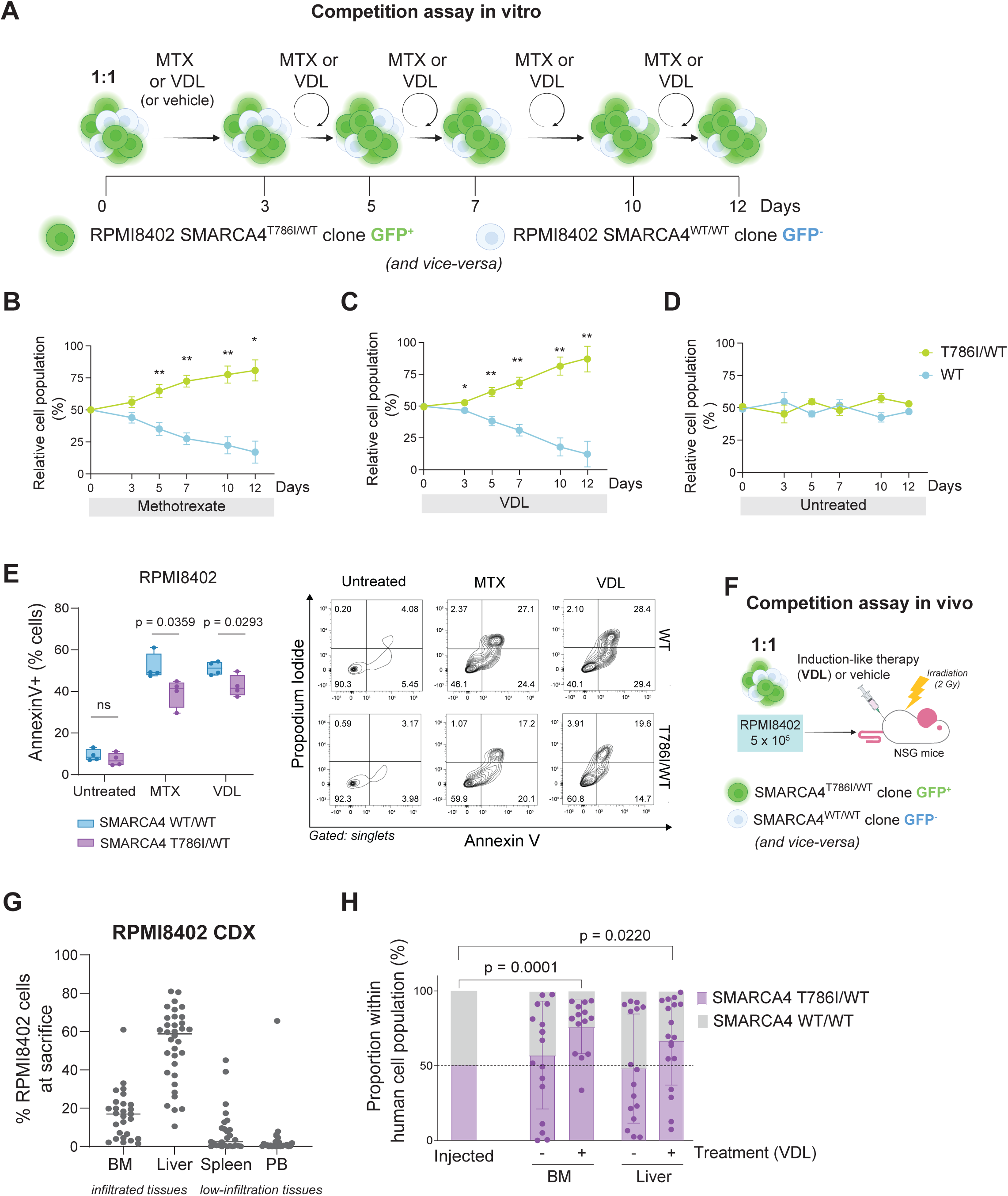
Relapse-associated SMARCA4 mutations confer a competitive advantage under chemotherapy pressure. **A.** Experimental design of in vitro competition assays performed using RPMI8402 WT and SMARCA4 T786I/WT clones. **B-D.** Relative abundance of RPMI8402 WT and SMARCA4 T786I/WT populations during MTX treatment (**B**), VDL treatment (**C**), or untreated culture (**D**). Four independent 1:1 mixes were analyzed: each of three individual GFP-labeled SMARCA4 T786I/WT clones competed against an unlabeled pool of WT clones, plus the reciprocal mix of a GFP-labeled pool of WT clones competed against an unlabeled pool of SMARCA4 T786I/WT clones (shown individually in Figure S2A-B). For each mix, values at each time point were calculated as the mean of three independent experiments; data represent the mean ± SD of the four mixes. **E.** Apoptosis analysis following 72 h treatment with MTX or VDL of RPMI8402 WT and SMARCA4 T786I/WT clones. Left, quantification of Annexin V-positive cells. Each dot represents an independent isogenic clone. Values for each clone were calculated as the mean of 2 independent experiments. Right, representative Annexin V/propidium iodide flow cytometry plots. **F.** Experimental design of in vivo competition assays performed using RPMI8402 WT and SMARCA4 T786I/WT clones. **G.** Percentage of RPMI8402 cells in bone marrow (BM), liver, spleen and peripheral blood (PB) at sacrifice. RPMI8402 cells identified as mouse CD45- human CD45+CD5+CD7+ by flow cytometry and gated on DAPI negative cells. Each dot represents an independent animal (pooled from 3 independent experiments). **H.** Relative contribution of WT and SMARCA4 T786I/WT RPMI8402 cells within the human leukemic compartment in the BM and liver of VDL-treated (+) and untreated (-) mice. “Injected” bar shows the 1:1 input ratio, and the dashed line indicates equal representation of both populations (50%) as at the start of the experiment. Each dot represents an independent animal. For panels G-H, data were pooled from 3 independent experiments, with both labeling strategies equally represented in each group. Only animals with leukemic engraftment above the predefined cut-off (2% in BM, 5% in liver) were included in panel H: BM, n = 17 untreated and n = 14 VDL-treated; liver, n = 16 untreated and n = 18 VDL-treated. For panels B-D and H, statistical significance was assessed at each time point using a two- tailed one-sample t-test against a theoretical value of 50%. For panel E, statistical significance between SMARCA4 WT and T786I/WT groups was determined using two-tailed unpaired t-tests. ****p < 0.0001, ***p < 0.001, **p < 0.01, *p < 0.05.

We next validated these findings in vivo using cell line-derived xenograft (CDX) models. Pools of isogenic RPMI8402 WT or SMARCA4-mutant (T786I/WT) clones (four independent clones per genotype) were mixed at a 1:1 ratio and transplanted into sublethally irradiated NSG mice (Figure 2F). To control for reporter-associated bias, we performed reciprocal labeling experiment, in which either the WT or mutant population carried the fluorescent reporter. Following leukemic engraftment, we randomized the mice into untreated or chemotherapy- treated groups receiving induction-like VDL^31^ (n = 8 per group; 4 per labeling strategy; 3 independent experiments). Animals were sacrificed upon development of overt leukemic disease, and we quantified the relative contribution of WT and SMARCA4-mutant populations across infiltrated tissues (Figure 2G). Under chemotherapy treatment, SMARCA4-mutant clones consistently outcompeted WT cells in the bone marrow, with a similar trend observed in the liver (Figure 2H) and in low-infiltration organs (Figure S2C). In contrast, untreated animals showed no significant enrichment of either population in infiltrated tissues, with considerable inter-individual variability. We observed similar results in the Jurkat model using an independent GFP/mCherry competition strategy (Figure S2D), where SMARCA4-mutant cells preferentially expanded within infiltrated organs following chemotherapy treatment (Figure S2F-S2G).

Together, these findings demonstrate that relapse-associated SMARCA4 mutations have a survival advantage under chemotherapy pressure in vivo.

### SMARCA4 mutations impair basal AP-1–associated enhancer accessibility and transcription of stress-responsive genes

Canonical SWI/SNF complexes play a central role in regulating chromatin accessibility and enhancer activity. To investigate whether relapse-associated SMARCA4 mutations reshape the chromatin landscape of T-ALL cells, we performed ATAC-seq in isogenic RPMI8402 SMARCA4-mutant (T786I/WT) and WT cells.

SMARCA4 mutation resulted in 3,734 differential accessible regions (DARs; Supplementary Data 1), with most changes (∼74%) corresponding to regions showing reduced accessibility relative to WT controls (DARs down: n = 2,797; DARs up: n = 937; Figure 3A), compatible with partial impairment of SWI/SNF-mediated chromatin remodeling. Similarly, SMARCA4 knockdown induced an even broader loss of chromatin accessibility (DARs down: n = 9,308; DARs up: n = 1,384; Figure S3A-C; Supplementary Data 1).

**Figure 3.**
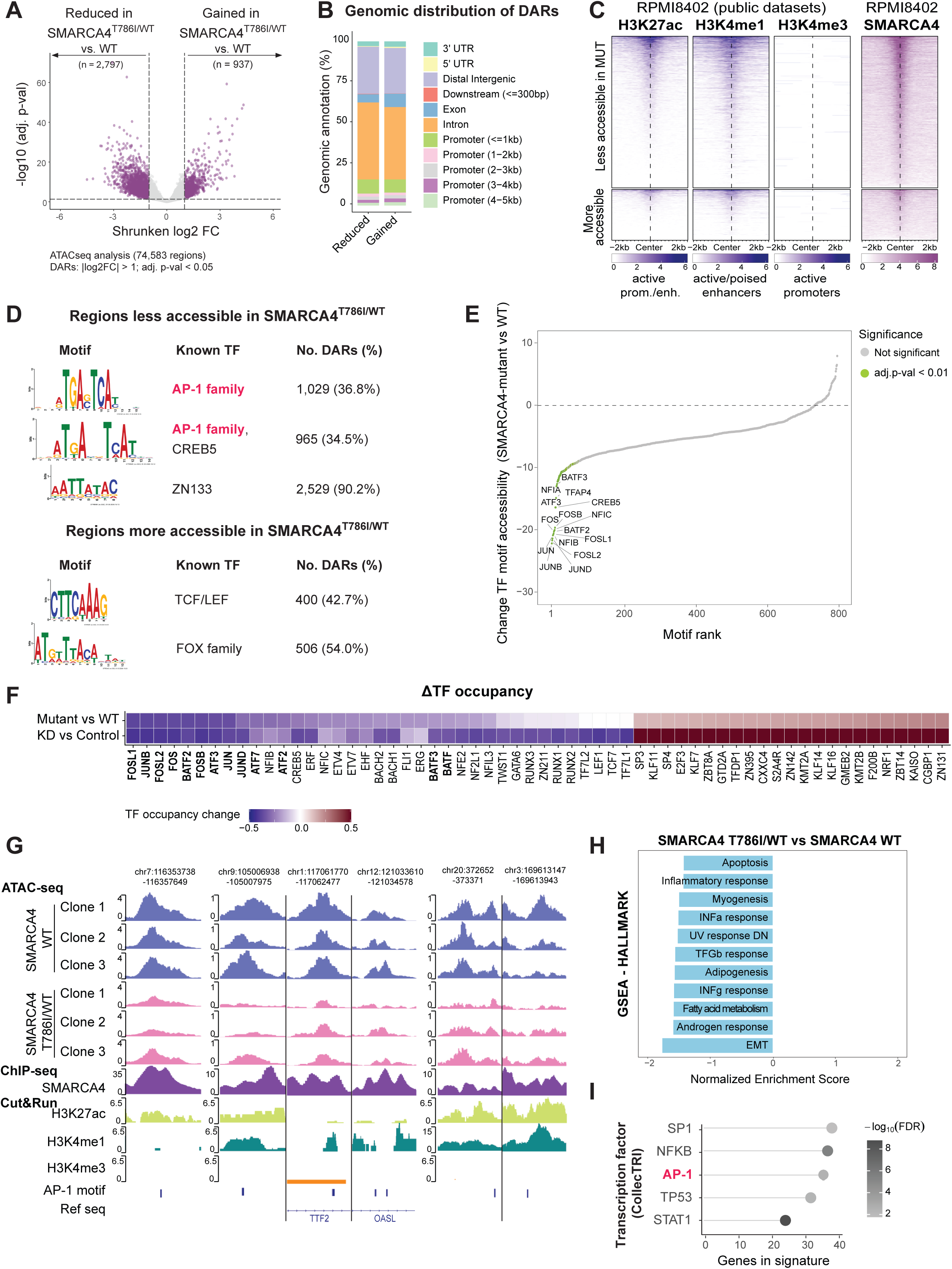
SMARCA4 mutations impair basal AP-1–associated enhancer accessibility and transcription of stress-responsive genes. **A.** Volcano plot showing differential chromatin accessibility between WT and SMARCA4 T786I/WT RPMI8402 cells under basal conditions, as determined by ATAC-seq. Differentially accessible regions (DARs) were defined as regions with an absolute shrunken log2FC > 1 and adjusted p-value (FDR) < 0.05. Total regions analyzed, n = 74,583; DARs with reduced accessibility, n = 2,797; DARs with increased accessibility, n = 937. **B.** Genomic annotation of regions showing reduced or increased accessibility in SMARCA4 T786I/WT cells, as defined in A. **C.** Heatmaps showing H3K27ac, H3K4me1, H3K4me3, and SMARCA4 occupancy in WT RPMI8402 cells at regions displaying reduced (top) or increased (bottom) accessibility in SMARCA4 T786I/WT cells. Signals are shown within a ±2 kb window centered on each region. Histone CUT&RUN-seq datasets were obtained from parental RPMI8402 cells (GSE231963; H3K27ac, GSM7306523; H3K4me1, GSM7306524; H3K4me3, GSM7306521). SMARCA4 ChIP-seq was performed under basal conditions in RPMI8402 cells, and tracks represent the average signal from three independent WT clones. **D.** Transcription factor motif enrichment analysis of regions displaying reduced (top) or increased (bottom) accessibility in SMARCA4-mutant cells. ATAC-seq regions without significant accessibility changes were used as the background set. Statistical significance thresholds were STREME E-value < 5 e^-02^ and Tomtom q-value < 5 e^-02^. For each motif, the indicated number and percentage correspond to DARs containing the respective motif. **E.** Ranking of transcription factor (TF) motif accessibility changes (chromVAR) in SMARCA4 T786I/WT versus WT RPMI8402 cells under basal conditions. Negative values indicate lower motif accessibility in SMARCA4 T786I/WT cells, and vice-versa. Motifs are ordered by the magnitude of the change. Motifs with a statistically significant change (adjusted p-value < 0.01) are highlighted in green and labeled. **F.** Differential transcription factor occupancy inferred by footprinting analysis in RPMI8402 SMARCA4 T786I/WT cells (versus WT) and in SMARCA4-knockdown cells (versus shControl). Statistical significance threshold: absolute ΔTF occupancy > 0.2 and p-value < 1 e^-100^. AP-1 family members are shown in bold. **G.** Representative genome browser (IGV) tracks showing reduced accessibility at AP-1- associated regulatory elements in independent RPMI8402 SMARCA4 T786I/WT clones. Corresponding SMARCA4 binding profiles and histone modification tracks from parental/WT RPMI8402 cells, together with AP-1 motif locations, are shown below. **H.** Bar plot showing gene set enrichment analysis (GSEA) of bulk RNA-seq data comparing SMARCA4 T786I/WT versus WT RPMI8402 cells under basal conditions, using the MSigDB Hallmark collection. Only statistically significant pathways are shown (adjusted p-value (FDR) < 0.1). Pathways are sorted in descending order of normalized enrichment score (NES). Full GSEA results are provided in Supplementary Data 5. **I.** Regulon overrepresentation analysis (CollecTRI) of leading-edge genes contributing to stress-response signatures altered in SMARCA4 T786I/WT cells (panel H). Top 5 statistically significant regulons are shown (adjusted p-value (FDR) < 0.1).

To determine whether the chromatin accessibility defects observed in SMARCA4-mutant cells were associated with altered SMARCA4 chromatin occupancy, we performed SMARCA4 ChIP- seq in WT and T786I/WT RPMI8402 cells under basal conditions. SMARCA4 occupancy profiles were largely comparable between genotypes across a consensus peak set (union of WT and mutant peaks, n = 20,567) (Figure S3E), and differential binding analysis identified only 55 significantly differential peaks (Supplementary Data 2). The T786I/WT mutation therefore impairs chromatin accessibility without substantially altering overall SMARCA4 chromatin occupancy.

Regions with impaired accessibility in SMARCA4-mutant (Figure 3B) or SMARCA4-depleted (Figure S3D) cells localized primarily to distal and intronic regions and displayed chromatin features characteristic of active enhancers, including high H3K27ac and H3K4me1 signals together with absence of H3K4me3 (Figure 3C). SMARCA4 ChIP-seq signal also accumulated around these sites (Figure 3C), supporting a direct role for SMARCA4 in maintaining accessibility at these regulatory regions. These data indicate that relapse-associated SMARCA4 mutations impair accessibility preferentially at enhancer-associated regulatory regions.

Regions losing accessibility in both SMARCA4-mutant and SMARCA4-depleted cells showed significant enrichment for AP-1 family motifs, including JUN, FOS, and ATF family members (Figure 3D, S3F; Supplementary Data 3). Accordingly, AP-1 family members ranked among the motifs with the greatest loss of accessibility in SMARCA4-mutant cells (Figure 3E). Footprinting analysis further predicted globally reduced occupancy across multiple AP-1 family members relative to WT controls (Figure 3F, S3G), suggesting impaired AP-1 engagement at chromatin. Notably, reduced AP-1 accessibility represented a shared feature of both SMARCA4 mutation and acute SMARCA4 depletion. Representative loci showed reproducible accessibility defects across independent SMARCA4-mutant clones at AP-1-associated regulatory regions (n = 3 clones per genotype, Figure 3G). Altogether, these data indicate that an intact SMARCA4 ATPase domain is required to maintain accessibility at a subset of AP-1- associated enhancer-like regions in T-ALL cells.

We next asked whether the changes in chromatin accessibility observed in basal SMARCA4 T786I/WT cells were accompanied by transcriptional dysregulation. RNA-seq analysis of isogenic SMARCA4-mutant and WT RPMI8402 cells revealed predominant transcriptional repression in SMARCA4-mutant cells (543 downregulated and 283 upregulated differentially expressed genes, DEGs; Supplementary Data 4), in agreement with the established role of SWI/SNF complexes in transcriptional activation^32,33^.

Despite the absence of overt phenotypic differences to WT under basal conditions, genes downregulated in SMARCA4-mutant cells were enriched for transcriptional programs linked to cellular stress responses, including apoptosis, inflammatory signaling or epithelial-to- mesenchymal transition, while no significantly enriched HALLMARK programs were observed among upregulated genes (GSEA adjusted p-value (FDR) < 0.1; Supplementary Data 5; Figure 3H).

To infer upstream regulators associated with these transcriptional changes, we performed regulon overrepresentation analysis against the CollecTRI database (a curated collection of TFs and their target genes^34^) on leading edge genes contributing to the enrichment in stress- related pathways (Supplementary Data 6). This analysis identified enrichment for AP-1- associated transcriptional programs among genes downregulated in SMARCA4-mutant cells, together with other stress-responsive regulators such as NFκB, TP53 and STAT1 (Figure 3I).

Thus, relapse-associated SMARCA4 mutations establish a pre-existing transcriptional state characterized by reduced expression of stress-response genes, including AP-1 targets, prior to chemotherapy exposure.

### SMARCA4-mutant cells fail to remodel AP-1-associated regions in response to chemotherapy

AP-1 transcription factors mediate rapid stress-induced enhancer activation and coordinate early transcriptional responses to stimuli^35–37^. To determine how relapse-associated SMARCA4 mutations affect these early chromatin responses to chemotherapy, we performed ATAC-seq in WT and SMARCA4-mutant (T786I/WT) RPMI8402 cells after short-term MTX exposure (24 h, IC50).

WT cells underwent chromatin remodeling in response to MTX, predominantly by increasing accessibility (1,426 regions gained accessibility and 657 regions lost accessibility; Figure 4A). In contrast, SMARCA4-mutant cells displayed a markedly attenuated response, with substantially fewer DARs detected after treatment (511 DARs up, 8 DARs down; Figure 4A). Most treatment-responsive regions in mutant cells were also remodeled in WT cells and changed in the same direction (Figure 4B, S4A), indicating that SMARCA4 mutation does not establish an alternative chromatin response program but rather reduces the magnitude of the normal response (Figure 4B,C). Principal component analysis (PCA) further supported this pattern, as SMARCA4-mutant cells remained closer to their basal chromatin state after treatment relative to WT controls (Figure S4B).

**Figure 4.**
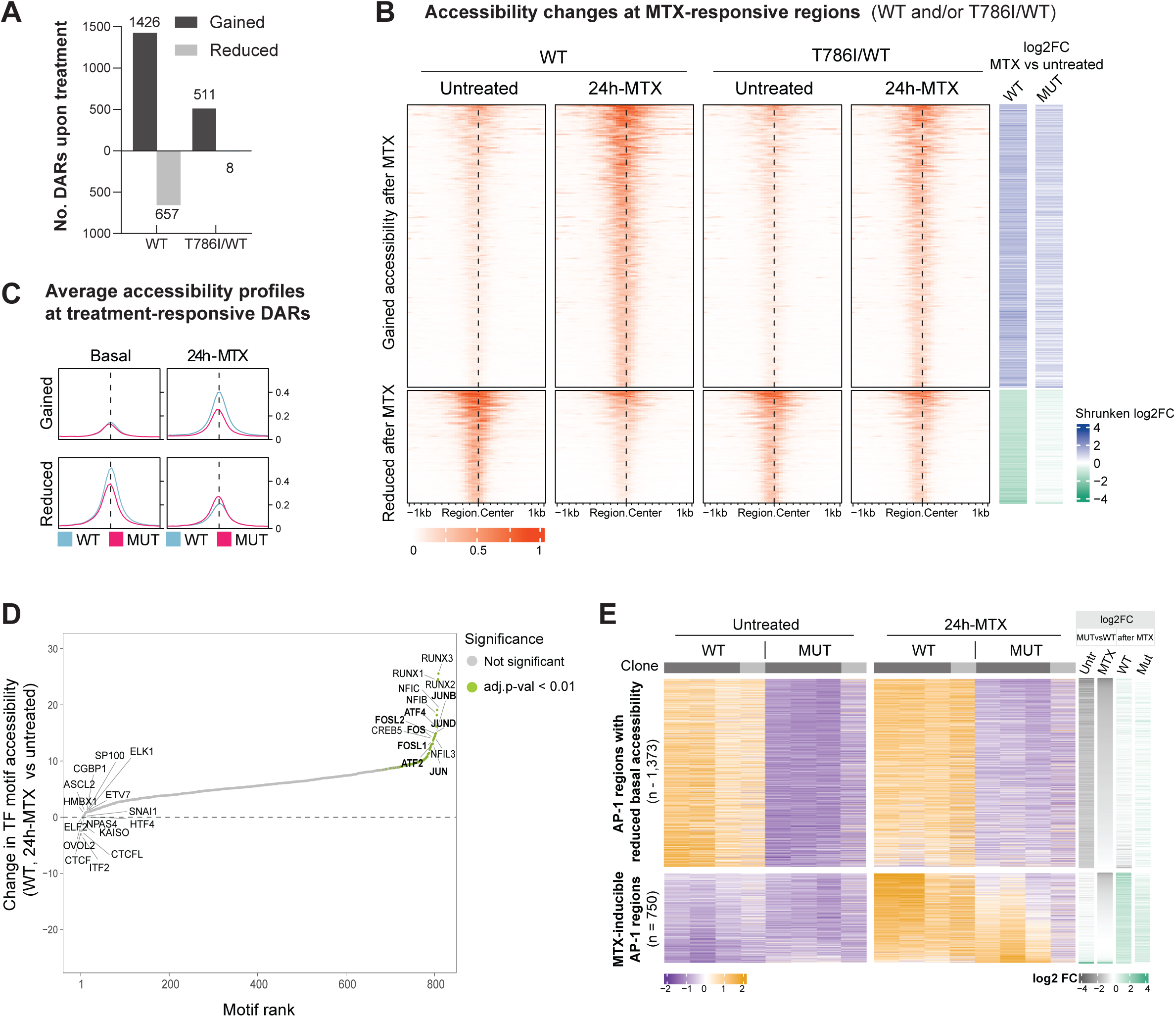
SMARCA4-mutant cells fail to remodel AP-1-associated chromatin in response to chemotherapy. **A.** Number of differentially accessible regions (DARs) gaining or losing accessibility after MTX treatment (24 h, 5 nM) in WT and SMARCA4 T786I/WT RPMI8402 cells. **B.** Heatmaps showing chromatin accessibility changes across treatment-responsive regions in WT and SMARCA4 T786I/WT RPMI8402 cells. Regions with gained accessibility (top, n = 1,652) and regions with reduced accessibility (bottom, n = 658) in either genotype after 24h MTX are represented. ATAC-seq signal represents the average of three biological replicates per experimental group. Regions are ordered by decreasing accessibility in untreated WT cells. Side annotations indicate the log2 fold change following MTX treatment relative to untreated controls in WT or SMARCA4 T786I/WT cells. **C.** Aggregate accessibility profiles illustrating chromatin remodeling dynamics following MTX treatment in WT and SMARCA4 T786I/WT cells. ATAC-seq signal represents the average of three biological replicates per experimental group. **D.** Ranking of TF motif accessibility changes (chromVAR) in WT RPMI8402 cells, comparing untreated and 24 h MTX-treated conditions. Positive values indicate higher motif accessibility after MTX treatment, and vice-versa. Motifs are ordered by the magnitude of the change. Motifs with a statistically significant change (adjusted p-value < 0.01) are highlighted in green and labeled. **E.** Heatmaps showing accessibility of two AP-1-associated enhancer classes before and after MTX treatment (24 h, 5 nM). Top, AP-1-associated regions displaying reduced basal accessibility in SMARCA4 T786I/WT cells (n = 1,373). Bottom, AP-1-associated regions undergoing inducible chromatin opening following MTX treatment (n = 750). Side annotations indicate the log2 fold change between SMARCA4 T786I/WT and WT cells under basal conditions (left) and the log2 fold change following MTX treatment relative to untreated controls (right). Heatmaps include three biological replicates from one WT and one SMARCA4 T786I/WT clone used for differential accessibility analyses, together with one additional WT and one SMARCA4 T786I/WT clone used for validation. Statistical thresholds used for differential accessibility analysis are indicated in Methods.

Analysis of motif accessibility showed that AP-1 family members were among the motifs with the largest increases after MTX exposure in WT cells, together with RUNX and NFI factors (Figure 4D); whereas reductions were smaller and showed no dominant motif (Figure 4D, S4C; Supplementary Data 3).

We next examined the behavior of AP-1-associated regulatory elements during treatment. AP- 1 associated regions with reduced accessibility in SMARCA4-mutant cells at baseline largely retained this differential state after treatment across independent clones (n = 1,373 regions, Figure 4E), indicating that pre-existing accessibility defects persist during chemotherapy treatment.

In addition, a distinct subset of AP-1–associated regions underwent inducible chromatin opening in WT cells upon treatment (n = 750, Figure 4E). These regions showed comparable accessibility between genotypes at baseline and overlapped only marginally with the AP-1- associated regions that are less accessible in untreated SMARCA4-mutant cells (n = 74 regions, 9.9%). Following treatment, however, these regions underwent substantially stronger remodeling in WT cells, whereas mutant cells showed little or only partial accessibility gains (Figure 4E). As a result, mutant cells failed to fully establish the AP-1 chromatin landscape normally acquired during chemotherapy stress, revealing both persistent basal defects and impaired inducible enhancer remodeling.

### SMARCA4-mutant cells display attenuated stress-response programs and retain fitness- associated pathways during chemotherapy exposure

We then investigated whether the impaired remodeling at AP-1 elements influenced transcriptional responses to MTX. RNA-seq analysis following acute MTX treatment (24 h, IC50) revealed robust transcriptional reprogramming in WT cells (upregulated DEGs: n = 1,809; downregulated DEGs: n = 513) (Figure 5A, Supplementary Data 4). This response was characterized by induction of stress-responsive programs including inflammatory response, apoptosis, TNFα/NFκB signaling, IL2/STAT5 signaling, p53 pathway activation and epithelial- to-mesenchymal transition (EMT), together with repression of pathways linked to cellular fitness and proliferation, such as MYC targets, E2F targets, G2M checkpoint and oxidative phosphorylation (Figure 5B, Supplementary Data 5). These changes are characteristic of the acute transcriptional response of tumor cells to cytotoxic stress^38,39^.

**Figure 5.**
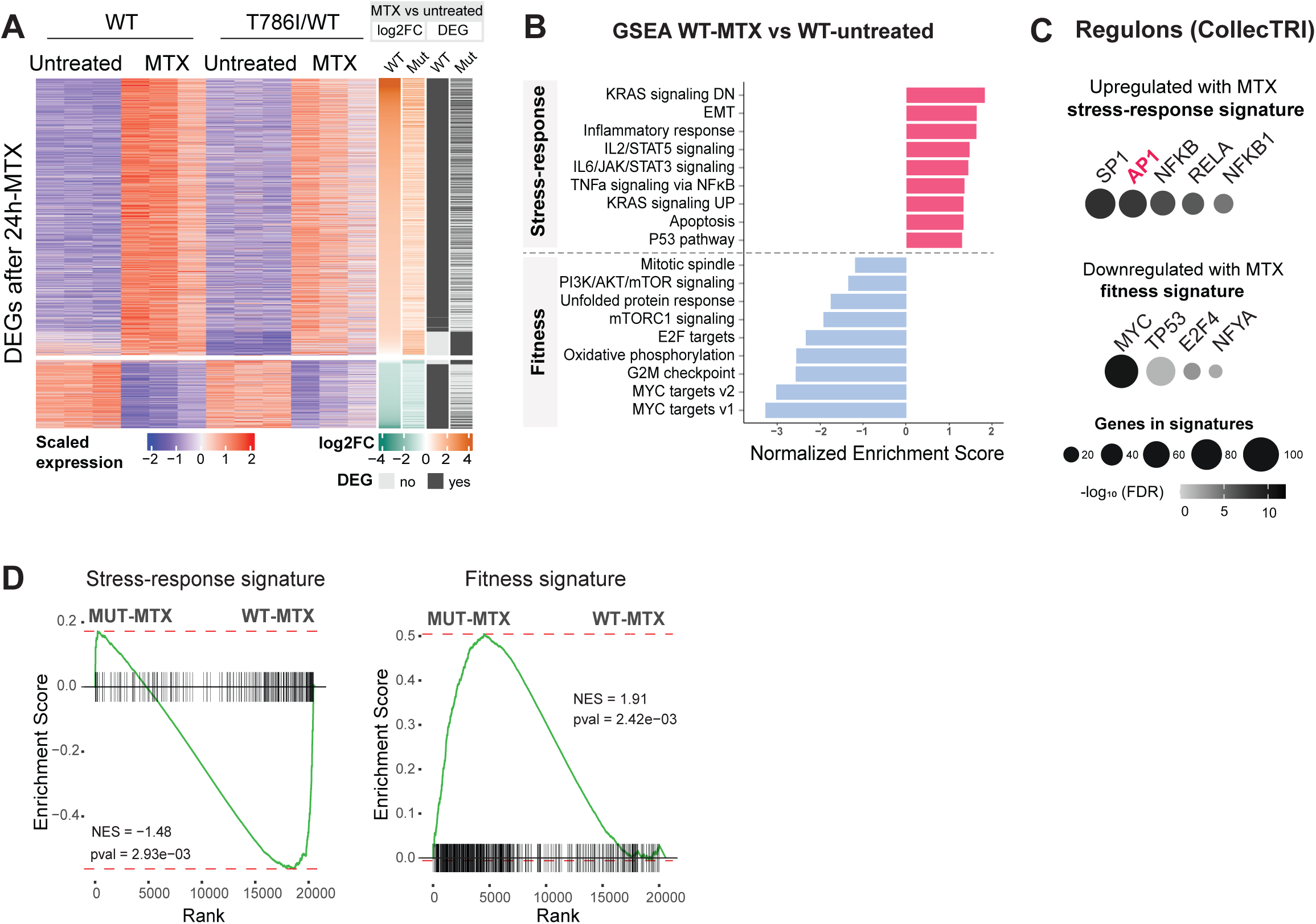
SMARCA4-mutant cells display attenuated stress-response programs and retain fitness-associated pathways during chemotherapy exposure. **A.** Expression heatmap of genes differentially regulated by MTX treatment (24 h, 5 nM) in WT and SMARCA4 T786I/WT RPMI8402 cells (WT, n = 1,809 upregulated and n = 513 downregulated; SMARCA4 T786I/WT, n = 920 upregulated and n = 149 downregulated). Columns represent WT and SMARCA4 T786I/WT cells under untreated and MTX-treated conditions (n = 3 independent replicates). Side annotations indicate the fold change and differential expression status for the MTX versus untreated comparison within each genotype. **B.** Bar plot showing normalized enrichment scores (NES) from pre-ranked GSEA of the MTX- treated versus untreated comparison in WT RPMI8402 cells, using the MSigDB Hallmark collection. Statistically significant pathways (adjusted p-value < 0.1) representative of stress- response and fitness-associated programs are shown, sorted by descending NES. Full GSEA results are provided in Supplementary Data 5. **C.** Regulon overrepresentation analysis (CollecTRI) of leading-edge genes contributing to MTX-responsive signatures. Only statistically significant regulons are shown (adjusted p-value (FDR) < 0.1). **D.** GSEA plots showing downregulation of stress-response signatures (left) and upregulation of fitness-associated signatures (right) in MTX-treated SMARCA4 T786I/WT compared with MTX-treated WT controls. The corresponding signatures were derived from leading-edge genes contributing to stress-response or fitness-associated pathways identified in panel B. The complete gene lists are provided in Supplementary Data 6. Statistical thresholds used for differential expression and functional analysis are indicated in Methods.

To identify the regulatory networks associated with this response, we analyzed the leading- edge genes contributing to the enrichment of stress-response pathways induced in WT cells (Figure 5C, Supplementary Data 6). Regulon overrepresentation analysis identified AP-1 family transcription factors, together with NFκB- and SP1-associated networks, among the most significantly enriched upstream regulators of these treatment-responsive genes. Conversely, genes contributing to pathways repressed by MTX were enriched for MYC- and E2F-associated transcriptional programs (Figure 5C).

Consistent with the attenuated chromatin remodeling, SMARCA4-mutant cells mounted a qualitatively similar but quantitatively attenuated transcriptional response to chemotherapy. MTX induced fewer transcriptional changes in mutant cells (upregulated DEGs: n = 920; downregulated DEGs: n = 149; Supplementary Data 4), and these were largely a subset of the genes responding in WT cells (n = 860 of 1,609; Figure 5A). Among the genes responding to MTX in both genotypes, fold changes were consistently smaller in mutant cells (Figure 5A). As a result, MTX-treated SMARCA4-mutant cells failed to fully activate stress-response genes, with relative preservation of fitness-associated transcriptional programs (Figure 5D). These findings indicate that relapse-associated SMARCA4 mutation impairs the establishment of the transcriptional state normally triggered by acute chemotherapy stress, including reduced activation of AP-1-associated stress-response networks.

### Reduced AP-1 chromatin engagement in SMARCA4-mutant cells accompanies chemotherapy tolerance

Our analyses showed that SMARCA4 mutation reduces accessibility at AP-1-associated regulatory regions (Figure 3) and inferred that the stress-responsive pathways downregulated in mutant cells are enriched for AP-1 targets (Figure 3I). To determine whether AP-1 occupancy is also altered, we performed ChIP-seq with c-Jun antibody in SMARCA4 WT and T786I/WT RPMI8402 cells under basal conditions. At the AP-1 associated regions showing reduced basal accessibility in mutant cells (n = 1,373, Figure 4E top), c-Jun occupancy was markedly reduced in SMARCA4 T786I/WT cells compared to WT (Figure 6A), while expression of AP-1 family members remained similar (Figure S5A). To extend this observation beyond the sequenced clones, we performed c-Jun ChIP-qPCR at three of these regions, together with an irrelevant genomic region as background control, in three independent clones for each genotype (Figure 6B). c-Jun enrichment was reduced at all three regions in SMARCA4 T786I/WT clones. This is consistent with the reduced AP-1 occupancy inferred from footprinting analyses (Figure 3F).

**Figure 6.**
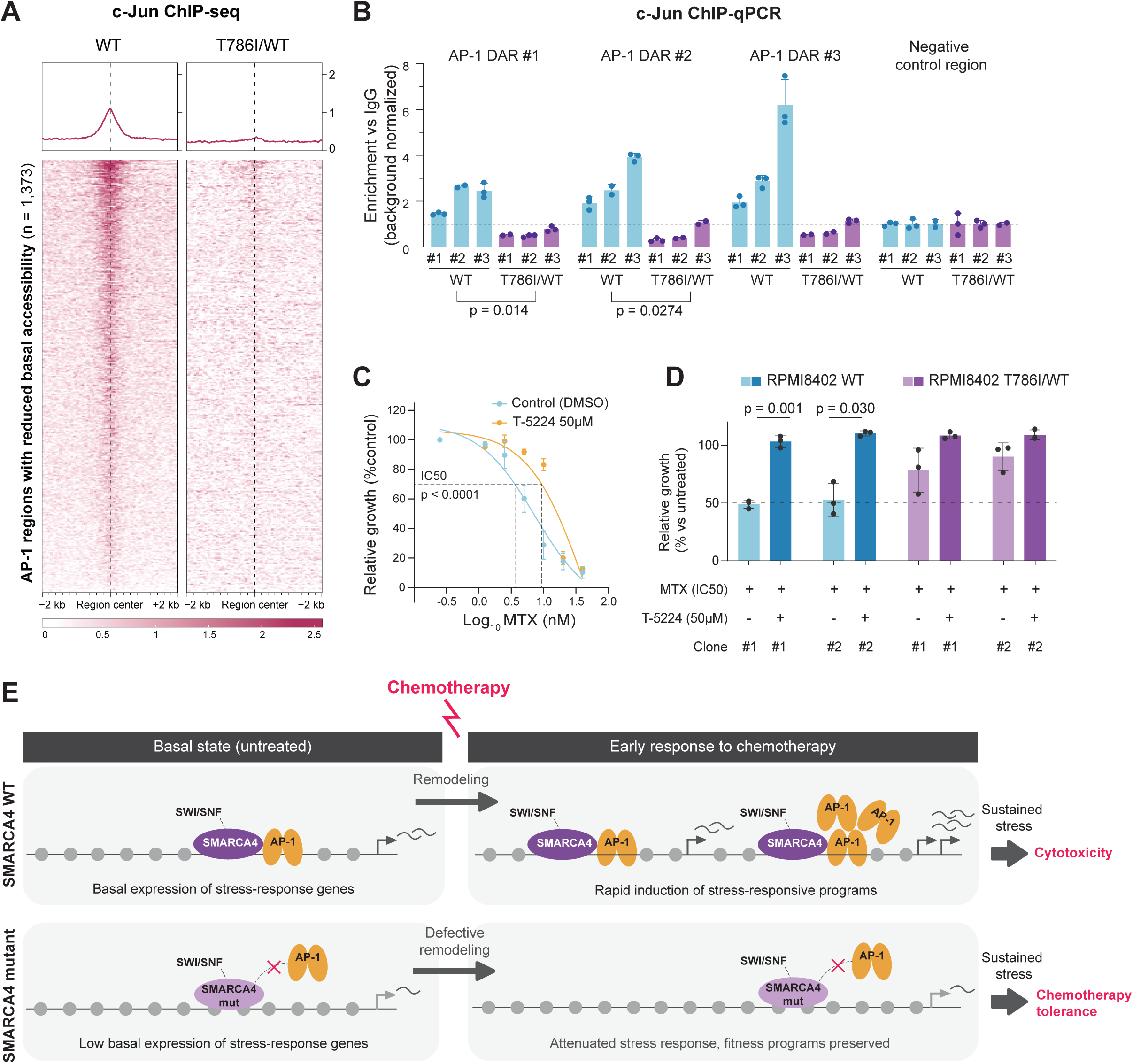
Reduced AP-1 chromatin engagement in SMARCA4-mutant cells accompanies chemotherapy tolerance. **A.** c-Jun ChIP-seq occupancy profiles in WT and SMARCA4 T786I/WT RPMI8402 cells under basal conditions, at AP-1-associated regions displaying reduced basal accessibility in SMARCA4 T786I/WT cells (n = 1,373). Signals are shown within a ±2 kb window centered on each region. c-Jun ChIP-seq was performed in the same WT and SMARCA4 T786I/WT clone used for differential accessibility and expression analyses. **B.** c**-**Jun ChIP-qPCR at three AP-1-associated regions (AP-1 DAR 1: chr20:372652-373371; AP-1 DAR 2: chr7:116353738-116357649; AP-1 DAR 3: chr12:121033610-121034578), and at an irrelevant negative control region (chr8:78022725-78022857), in three independent WT and three independent SMARCA4 T786I/WT RPMI8402 clones (#1 sequenced clones, #2-#3, two additional independent clones). Enrichment is expressed relative to IgG and normalized to the irrelevant negative control region as background. Each dot represents a technical replicate; bars represent the mean ± SD. Statistical significance between SMARCA4 WT and T786I/WT groups was determined at each region using two-tailed unpaired t-tests, considering each clone an independent biological replicate. **C.** Sensitivity of parental RPMI8402 cells to MTX (72 h) in the presence of the c-Fos/c-Jun inhibitor T-5224 (50 µM) or vehicle (DMSO), shown as dose-response curves with the IC50 of vehicle-treated cells indicated. Data represent the mean ± SD of three independent experiments. IC50 values were compared between conditions using two-tailed unpaired t-tests. **D.** Relative growth of two independent WT and two independent SMARCA4 T786I/WT RPMI8402 clones treated with MTX (72h, WT IC50) alone or combined with T-5224 (50 µM). Growth is normalized to vehicle-treated cells of each clone; dashed line, 50%. Bars show mean ± SD of three independent experiments; each dot represents one experiment. MTX and MTX plus T-5224 were compared within each clone using two-tailed unpaired t-tests. **E.** Model representing chromatin remodeling at AP-1-associated enhancers in cells carrying SMARCA4 mutations. In WT cells, a subset of AP-1-associated enhancers is maintained in an accessible state prior to treatment, supporting basal expression of stress-response genes. Chemotherapy exposure triggers additional AP-1-dependent enhancer remodeling, resulting in rapid activation of stress-responsive transcriptional programs, growth arrest and apoptosis. In contrast, relapse-associated SMARCA4 mutations reduce accessibility at pre-existing AP-1- associated enhancers and impair inducible AP-1-dependent chromatin remodeling following chemotherapy exposure. Consequently, SMARCA4-mutant cells exhibit attenuated activation of stress-response programs, relative preservation of fitness-associated pathways, reduced apoptosis and increased chemotherapy tolerance.

Finally, to test whether reduced AP-1 activity is responsible for chemotherapy tolerance in RPMI8402 T-ALL cells, we treated parental cells with T-5224, a selective inhibitor of c-Fos/c- Jun DNA-binding activity. AP-1 inhibition reduced MTX sensitivity in these cells, partially recapitulating the phenotype observed in SMARCA4-mutant cells (Figure 6C, S5B). At the concentration used, T-5224 alone did not affect cell growth (Figure S5C). AP-1 inhibition also promoted growth under MTX in WT clones, and had no significant effect in SMARCA4-mutant clones with already compromised AP-1 activity, which were already largely resistant to MTX alone (Figure 6D). These findings support a functional contribution of impaired AP-1 activity to the chemotherapy-tolerant state associated with relapse-related SMARCA4 mutations.

## DISCUSSION

In this study, we address the role of the SWI/SNF complex in the capacity of leukemic T-ALL cells to respond to chemotherapy. By following up on the previous observation that SMARCA4 mutations are found in about 10% of relapse T-ALL patients, we investigated the cause-effect of these mutations in the treatment response. Isogenic T-ALL cell lines engineered with the specific relapse-associated mutations in SMARCA4 ATPase domain showed more resistance to general chemotherapy agents than normal WT cells. The fact that we tested different mutations in 3 isogenic cell line models and 4 different types of chemotherapy treatments, including MTX and a cocktail of VDL, strongly suggests that this is a general mechanism at least in T-ALL, but that could also be operating in other cell types.

Acute depletion or pharmacological inhibition of SMARCA4 profoundly impairs leukemia maintenance in AML and T-ALL^13,21–23^, highlighting the essential role of SWI/SNF activity in these malignancies. In contrast, the heterozygous relapse-associated mutations studied here had minimal effects on basal proliferation or viability despite conferring broad resistance to multiple chemotherapeutic agents. These observations indicate that cells bearing relapse- associated mutations do not phenocopy all the traits of SMARCA4 loss. Instead, they share the chromatin remodeling effects without compromising leukemia maintenance.

We found that chromatin accessibility was reduced at AP-1-associated enhancer-like regions in SMARCA4-mutant cells under basal conditions. Representative regions also showed reduced JUN occupancy, despite unchanged expression of AP-1 family members. AP-1 factors recruit SWI/SNF to locally open stress-responsive enhancers^36,40^, which in turn further supports and stabilizes AP-1 binding. We propose that impaired remodeling by mutant SMARCA4 destabilizes this cooperation, so that reduced accessibility and reduced AP-1 occupancy reinforce each other at these elements. Consistent with a functional contribution of this defect, pharmacological inhibition of AP-1 DNA binding reduced MTX sensitivity in WT cells.

Following MTX treatment, these accessibility defects persisted, with substantially less chromatin remodeling than in WT cells, indicating that SMARCA4 ATPase activity is required to mount an effective chromatin response to chemotherapy. As a result, we found a weaker induction of inflammatory, apoptotic, and other stress-response programs, whereas programs associated with cellular fitness were comparatively preserved. We propose that mutant cells enter chemotherapy exposure from a pre-existing state in which AP-1-associated enhancers are both less accessible and less occupied, limiting the basal expression of stress-related genes and the capacity to amplify these programs under stress. This preconfigured chromatin state restricts activation of cytotoxic stress responses and thereby confers a selective advantage under chemotherapy pressure (Figure 6E).

This concept closely parallels the phenomenon of phenotypic inertia recently described by Loukas et al.^38^, in which disruption of diverse chromatin regulators (including members of the SWI/SNF complex) limits stress-induced transcriptional reprogramming and preserve cellular fitness. Rather than promoting constitutive resistance programs, perturbation of chromatin regulators reduces the magnitude of the acute stress response, thus increasing the probability of survival under adverse conditions. Our findings extend this principle to chemotherapy resistance and suggest that AP-1-dependent diminished chromatin responsiveness may represent a convergent mechanism through which diverse chromatin alterations promote treatment tolerance.

AP-1 transcription factors are key regulators of inducible enhancers activated by environmental and cellular stress, where they cooperate with SWI/SNF complexes to establish and remodel accessible chromatin^36,40^. Many of these enhancers are already occupied by AP-1 and maintained in an accessible state before stimulation, allowing rapid transcriptional activation following stress^35^. Our findings fit this framework by suggesting that relapse-associated SMARCA4 mutations impair the SWI/SNF-dependent remodeling steps required for AP-1 enhancer maintenance and activation. This provides a mechanistic explanation for the reduced chromatin accessibility and attenuated stress-induced transcription observed in mutant cells.

Recent studies have also placed AP-1-dependent enhancer remodeling at the center of adaptive responses to cancer therapy, where repeated exposure to increasing drug concentrations progressively rewires AP-1-associated enhancer landscapes and enables tumor cells to explore alternative transcriptional states until stable resistant phenotypes emerge^41–43^. These studies further suggest that survival under initial high-dose therapy depends primarily on pre-existing cellular states rather than gradual adaptation. Our experimental model captures this acute phase of chemotherapy, which more closely reflects the clinical treatment of T-ALL. In this context, relapse-associated SMARCA4 mutations may establish a chromatin state that already shares key features with the adapted endpoint described in these studies, including attenuated stress-associated transcription and reduced chromatin responsiveness following chemotherapy. Thus, impaired SWI/SNF remodeling may provide a genetic shortcut toward a stress-tolerant state, reducing the need for the extensive enhancer reprogramming normally required to achieve chemotherapy resistance.

The results presented in this work raise several important questions. A key priority will be to determine how relapse-associated ATPase-domain mutations affect the activity of distinct SWI/SNF assemblies and whether mutant and WT SMARCA4-containing complexes make different contributions to chromatin remodeling in the heterozygous state. It will also be important to establish whether this mechanism extends to other leukemias and malignancies. Beyond its biological implications, the altered regulatory state imposed by SMARCA4 mutations is likely to create therapeutic vulnerabilities that remain to be identified. Defining these dependencies may reveal new strategies to overcome chemotherapy tolerance in relapsed disease.

Overall, our findings establish SMARCA4 ATPase activity as a critical regulator of the chromatin response to chemotherapy and identify impaired chromatin responsiveness as a mechanism of treatment tolerance in T-ALL. More broadly, they suggest that the dynamic response of the cancer epigenome to therapeutic stress is a critical determinant of treatment outcome. Together, these results identify chromatin remodeling pathways as potential therapeutic vulnerability in resistant cells, yet underscore the need for caution when using chromatin modifier agents in combination with chemotherapy, as such combination may unexpectedly promote resistance.

## MATERIALS AND METHODS

### CELL CULTURES AND REAGENTS

#### Cell lines

The T-cell acute lymphoblastic leukemia (T-ALL) cell lines RPMI8402 (DSMZ ACC- 290) and Jurkat (DSMZ ACC-282) were cultured in RPMI-1640 medium (Sigma, R8758; containing L-glutamine) supplemented with 10% fetal bovine serum (Gibco, A5256701), 1% penicillin-streptomycin (Sigma, P4333), and 1% sodium pyruvate (Sigma, S8636). HEK293T cells (ATCC CRL-3216) were cultured in Dulbecco’s Modified Eagle Medium (DMEM; Sigma, D6546) supplemented with 10% fetal bovine serum (Gibco, A5256701), 1% penicillin- streptomycin (Sigma, P4333), 1% L-glutamine (Sigma, G7513), and 1% sodium pyruvate (Sigma, S8636). All cell lines were maintained at 37°C in a humidified atmosphere containing 5% CO₂ and were routinely tested for Mycoplasma contamination. Mycoplasma-free status was confirmed by PCR before each experiment.

#### Reagents and drug treatments

Clinical-grade methotrexate (25 mg/mL), vincristine (1 mg/mL), dexamethasone (4 mg/mL), doxorubicin (2 mg/mL), and etoposide (20 mg/mL) were kindly provided by the Hospital del Mar Pharmacy Department (Barcelona, Spain). L- asparaginase (MedChemExpress, HY-P1923) stock solution was prepared in 0.2 M Tris-HCl (pH 7.0). AP-1 inhibitor T-5224 (MedChemExpress, HY-12270) stock solution was prepared in dimethyl sulfoxide (DMSO; Sigma-Aldrich). Unless otherwise indicated, compounds were diluted to their final working concentrations immediately before use in culture media.

#### MOUSE MODELS

All animal procedures were performed under specific pathogen-free (SPF) conditions, in compliance with the institutional and governmental guidelines for the care and use of laboratory animals, as established by the Animal Care Committee of the Generalitat de Catalunya. All experimental protocols were approved by the Committee for Animal Experimentation of the Barcelona Biomedical Research Park (PRBB).

### GENERATION OF ISOGENIC SMARCA4 HETEROZYGOUS MUTANT T-ALL CELL LINES

#### Design of CRISPR reagents

sgRNAs were designed using Benchling to generate double- strand breaks within less than 10 bp of the desired SMARCA4 variants (T786I and R1192H), to maximize homology-directed repair (HDR) efficiency. Repair templates consisted of 100-nt single-stranded donor oligonucleotides carrying the desired nucleotide substitution centered within the donor sequence together with synonymous mutations disrupting the protospacer adjacent motif (PAM) to prevent Cas9 re-cutting following HDR. Additional silent nucleotide substitutions introducing de novo restriction enzyme recognition sites were incorporated to facilitate rapid identification of edited clones.

#### CRISPR/Cas9 genome editing

Isogenic SMARCA4-mutant T-ALL cell lines were generated by CRISPR/Cas9-mediated homology-directed repair (HDR). CRISPR ribonucleoprotein (RNP) complexes were assembled using 3 μL of 62 μM Alt-R S.p. HiFi Cas9 Nuclease V3 (Integrated DNA Technologies, IDT) and 5 μL of 44 μM gene-specific synthetic sgRNAs (Alt-R CRISPR-Cas9 sgRNA, IDT). Repair templates consisted of 100-nt single-stranded donor oligonucleotides (10 μL of 44 μM per reaction; Ultramer DNA Oligos, IDT) carrying the desired nucleotide substitution together with synonymous mutations disrupting the protospacer adjacent motif (PAM) to prevent Cas9 re-cutting after HDR. Donor oligonucleotides also contained an additional silent nucleotide substitution introducing a de novo restriction enzyme recognition site (AvaI for the T786I donor and SacI or ApaI for the R1192H donors), to facilitate rapid screening of edited clones. To generate heterozygous SMARCA4-mutant RPMI8402 clones, WT and mutant donor oligonucleotides were mixed at a 1:1 ratio before transfection. To revert the endogenous R1192H mutation in Jurkat cells, only the WT donor was used. RNP complexes and donor templates were delivered into 1 × 10^6^ cells resuspended in 92 μL of Buffer R using the Neon Transfection System (Thermo Fisher Scientific) with 100-μL electroporation tips at 1350 V, 10 ms, and 3 pulses. Following electroporation, cells were recovered for approximately 40 h in antibiotic-free medium containing 1 μM Alt-R HDR Enhancer V2 before analysis of editing efficiency and single-cell cloning.

#### Isolation of clonal cell lines

Approximately 40 h after electroporation, single cells were isolated by limiting dilution (0.75 cells per well) or fluorescence-activated cell sorting (FACS) into 96-well plates containing a 3:1 mixture of fresh and conditioned culture medium. Clones were expanded for 2–3 weeks, with medium replenished every 2–3 days. Once clones reached sufficient density, we kept a fraction of each culture for continued expansion, and harvested the remaining cells for genomic DNA extraction and genotyping. Clones carrying the desired genotype were subsequently expanded and cryopreserved.

#### Genotyping by Sanger sequencing

Genomic DNA was extracted from expanded clones by colony PCR. The targeted SMARCA4 genomic regions were amplified using locus-specific primers, and PCR products were purified and subjected to Sanger sequencing. Sequencing chromatograms were analyzed to identify clones carrying the desired heterozygous mutation and to confirm the absence of undesired editing events. In parallel, total RNA was extracted and reverse transcribed, and the corresponding cDNA regions were amplified and sequenced to confirm the mutational status and expression of the mutant alleles at the mRNA level.

sgRNA sequences and donor oligonucleotides are listed in Appendix Table 1. PCR and Sanger sequencing primers are listed in Appendix Tables 2.

## LENTIVIRAL PRODUCTION

Lentiviral particles were produced in HEK293T cells (ATCC CRL-3216). Packaging plasmids (Addgene #8455 and #12259) were co-transfected with the plasmid of interest using polyethyleneimine (PEI): pLKO.1-Puro shRNA transfer plasmids for knockdown, or pCDH-EF1a-eFFly-eGFP (Addgene #104834) and pCDH-EF1a-nanoLUC- mCherry (generated in house) for fluorescent labelling. Viral supernatants were harvested, filtered through 0.45-µm membranes, concentrated overnight using Lenti-X Concentrator (Takara Bio, #631232), and pelleted by centrifugation at 1,500 × g for 45 min at 4 °C.

## SMARCA4 KNOCKDOWN BY SHRNA

RPMI8402 cells were transduced with lentiviral particles encoding SMARCA4-targeting shRNAs (MISSION® shRNA clone TRCN0000218544 (shSMARCA4 #1) or TRCN0000231101 (shSMARCA4 #2); Sigma-Aldrich) or a non-targeting scrambled control shRNA (SHC016; Sigma-Aldrich) in the presence of 10 μg/mL polybrene. Transduced cells were selected and maintained in 1.25 μg/mL puromycin. SMARCA4 knockdown was confirmed by RT-qPCR and WB analysis relative to shControl.

## WESTERN BLOT ANALYSIS

For cytoplasm/nuclear fractionation, cells were lysed in ice-cold Buffer A (10 mM Hepes, 1.5 mM MgCl_2_, 10 mM KCl, and 0.05% NP-40; pH 7.9) for 15 min on ice, followed by centrifugation 800 × *g* for10 min at 4°C. Supernatants were recovered and mixed with 6X Laemmli buffer (cytoplasmic fraction). The remaining pellet fraction was resuspended in 1X Laemmli buffer and sonicated 2X for 10 s with 10% amplitude (nuclear fraction). Protein samples were denatured at 95°C for 10 min, resolved by SDS-polyacrylamide gel electrophoresis (SDS-PAGE) and analyzed by WB using standard procedures. Antibodies used are listed in Appendix Table 3.

## RT-qPCR ANALYSIS

Total RNA was isolated using the RNeasy Mini Kit (QIAGEN), and cDNA was synthesized using the Transcriptor First Strand cDNA Synthesis Kit (Roche). Quantitative real-time PCR (RT-qPCR) was performed on a LightCycler 480 Instrument II (Roche) using LightCycler 480 SYBR Green I Master (Roche). Relative gene expression was normalized to the housekeeping genes *TBP* and *HPRT1* and calculated using the 2^−ΔΔCt^ method. Primer sequences are listed in Appendix Table 2.

## CELL PROLIFERATION AND VIABILITY ASSAYS

Cell proliferation and viability were assessed over the indicated culture period using a Countess II Automated Cell Counter (Thermo Fisher Scientific). At each time point, cell suspensions were mixed with Trypan Blue solution to determine the number of viable cells and the percentage of viable (Trypan Blue- negative) cells.

## CELL CYCLE ANALYSIS

Cell cycle distribution was determined by flow cytometry using Ki67- AF647 and DAPI intracellular stainings. Unsynchronized RPMI8402 cells were collected after 48 h culture under standard conditions. Fixed and permeabilized cells (FIX&Perm Cell permeabilization kit, Invitrogen GAS004) were stained with Ki67-AF647 (1/100, 20 min in the dark; clone B56, BD Bioscience 558615) and DAPI (5 μg/ml, 1 h in the dark). Samples were acquired on a LSR Fortessa Cytometer (BD Bioscience). Data analysis was performed using FlowJo v10.10.1 (BD Bioscience).

## APOPTOSIS ANALYSIS

Apoptosis was quantified by Annexin V/propidium iodide (PI) staining followed by flow cytometry. Four independent SMARCA4 WT and four independent SMARCA4-mutant (T786I/WT) clones were cultured for 72 h under basal conditions or treated with 5 nM methotrexate (MTX) or VDL (1 nM vincristine, 1 μM dexamethasone, and 1 × 10-5 U/mL L-asparaginase) before staining with APC-conjugated Annexin V and PI according to the manufacturer’s instructions. Samples were acquired on an LSRFortessa flow cytometer (BD Biosciences), and data were analyzed using FlowJo v10.10.1 (BD Biosciences). Two independent experiments were performed using four biological replicate clones per genotype.

## DRUG SENSITIVITY ASSAYS

T-ALL cell lines were seeded at 25,000 cells per well in 96- well plates and incubated with the indicated compounds and doses for 72 h. Single-agent concentration ranges are indicated in the corresponding figures. VDL was applied as a fixed combination of vincristine (0.1-10 nM), dexamethasone (0.1-10 µM) and L-asparaginase (1 × 10⁻⁷ to 1 × 10⁻² U/mL). Cell viability was assessed using the MTT assay (Sigma, M5655) according to the manufacturer’s instructions. Vehicle-treated cells (untreated condition) were used as the 100% viability reference for normalization. Dose–response curves were generated by nonlinear regression using a logistic model [log(inhibitor) vs. response] in GraphPad Prism v10.6.1 (GraphPad Software). Area under the curve (AUC) and IC50 values were calculated from the fitted curves. All treatments were performed in technical triplicates and repeated in at least three independent experiments.

For competition and apoptosis assays, cells were treated with 1 nM vincristine, 1 µM dexamethasone and 1 × 10⁻⁵ U/mL L-asparaginase, or 5 nM MTX corresponding to the IC50 in WT cells.

For AP-1 inhibition, parental RPMI8402 cells or isogenic WT and SMARCA4 T786I/WT clones were seeded as described above and treated for 72 h with the MTX concentration range used for single-agent drug sensitivity assays, in the presence of 50 µM T-5224 or DMSO. Cell viability was assessed by MTT, and dose-response curves, AUC and IC50 values were obtained as described above. Relative growth is expressed as a percentage of the corresponding vehicle-treated control for each condition.

### COMPETITION ASSAYS

#### Fluorescent labelling

Isogenic RPMI8402 and Jurkat SMARCA4 WT and mutant clones were transduced with lentiviral particles encoding eFFly-eGFP (Addgene #104834) or nanoLUC- mCherry (generated in house) as described above, and labeled populations were purified by FACS on the basis of GFP or mCherry expression.

#### In vitro competition assays

Labeled and unlabeled populations were mixed at a 1:1 ratio and seeded at 2.5 x 10^5^ cells/mL. For RPMI8402, each of three independent GFP-labeled SMARCA4 T786I/WT clones was competed against an equicellular pool of three unlabeled WT clones. To control for bias introduced by the labeling strategy, the reciprocal combination (unlabeled pool of T786I/WT clones versus a GFP-labeled pool of WT clones) was performed in parallel. Mixed cultures were left untreated or treated with 5 nM methotrexate or with VDL (1 nM vincristine, 1 µM dexamethasone and 1 × 10⁻⁵ U/mL L-asparaginase), corresponding to the IC50 of each treatment in WT cells. Medium and drug were refreshed every 2–3 days, adjusting cell density. The proportion of labeled and unlabeled cells was verified to be 50:50 at day 0 and quantified at each medium change on a BD LSR II cytometer. Data were analyzed with FlowJo v10.10.1 (BD Biosciences). Cultures were maintained until the total viable cell number fell below 1 × 10⁵. Each clone combination was assayed in three independent experiments.

For Jurkat, a pool of four GFP-labeled unedited (R1192H/WT) clones was mixed with a pool of four mCherry-labeled reverted (WT) clones, and the reciprocal labeling combination was performed in parallel. Treatment, sampling and analysis were as described above.

#### In vivo competition assays

NSG mice (strain: ANB//NOD.Cg-Prkdcscid Il2rgtm1Wjl/SzJ; 2- 4month-old males and females), were sublethally irradiated (2 Gy) and injected via the tail vein with 5 × 10⁵ RPMI8402 cells, or intratibially with 4 × 10⁵ Jurkat cells, consisting of a 1:1 mixture of SMARCA4-mutant and WT cells. For RPMI8402, mixtures comprised a pool of three GFP- labeled T786I/WT clones and a pool of three unlabeled WT clones, or the reciprocal labelling; for Jurkat, a pool of four GFP-labelled unedited (R1192H/WT) clones and a pool of four mCherry-labelled reverted (WT) clones, or the reciprocal. Eight animals were injected per mixture (n = 16 per experiment). Leukemic engraftment and evolution was monitored by in vivo bioluminescence imaging. Upon first detection of luciferase signal (within one week of injection), animals were randomized into two groups with comparable disease burden and balanced across labelling strategies: vehicle (PBS) or an induction-like chemotherapy combination (VDL)^31^ consisting of vincristine (0.15 mg kg⁻¹ day⁻¹, days 0 and 7), dexamethasone (5 mg kg⁻¹ day⁻¹, days 0, 5, 7 and 8) and L-asparaginase (1,000 IU kg⁻¹ day⁻¹, days 0, 5, 7 and 8). Drugs were diluted in sterile PBS and administered intraperitoneally. Disease progression was followed weekly by bioluminescence imaging and by flow cytometric analysis of peripheral blood. Animals were euthanized approximately three weeks after injection, when the first animals reached humane endpoints. Bone marrow, peripheral blood, spleen and liver were collected and analyzed on a BD LSR II cytometer. Leukemic cells were identified as mouse CD45⁻ human CD45⁺CD5⁺CD7⁺ within DAPI-negative events, and the relative contribution of GFP⁺ and GFP⁻ (or mCherry⁺) cells within the human compartment was quantified. Data shown are pooled from three independent experiments. Animals that died during the procedure and animals whose engraftment fell below the predefined cut-off (1% cells in peripheral blood, 2% of cells in bone marrow, 5% in liver and spleen) were excluded from the analysis.

### ASSAY FOR TRANSPOSASE-ACCESSIBLE CHROMATIN (ATAC)-SEQUENCING

#### Sample preparation

Isogenic RPMI8402 SMARCA4 WT and T786I/WT cells were seeded at 2.5 x 10^5^ cells/ml and cultured in complete RPMI1640 medium under basal conditions or following treatment with 5 nM methotrexate (MTX) for 24 h. After 24 h, viable cells (DAPI negative) were FACS-sorted and nuclei were isolated from 100,000 viable cells using the Omni- ATAC protocol^44,45^ with detergent-mediated lysis employing digitonin, Tween-20, and IGEPAL CA-630. Tagmentation was performed using the Nextera DNA Library Preparation Kit (Illumina) for 45 min at 37°C with agitation at 1,000 rpm. Following transposition, DNA was purified using the MinElute Reaction Cleanup Kit (QIAGEN) and submitted for library preparation and high- throughput sequencing.

Three independent cultures of one SMARCA4 WT clone and one T786I/WT clone were profiled under basal conditions and after 24 h MTX treatment and used for all differential chromatin accessibility analyses. Parallel RNA samples were collected from the same cultures for RNA sequencing. To assess the reproducibility of the identified accessibility changes across independently derived clones, two additional WT and T786I/WT clone pairs were profiled, including one clone pair under basal conditions only, and a second clone pair under both basal and MTX-treated conditions.

An independent ATAC-seq experiment was performed in RPMI8402 cells transduced with either shControl or shSMARCA4 under basal conditions, using three independent lentiviral transductions per condition.

#### Library preparation and sequencing

DNA quality control, library preparation, and sequencing were performed at the CRG Genomics Unit (PRBB, Barcelona, Spain). In each experiment, all libraries were prepared simultaneously according to the facility’s standard protocol and sequenced on an Illumina NovaSeq6000 platform (50 bp paired-end reads). Samples sequencing depth ranged between 74.6M and 91.6M reads or between 67.2M and 128.1M reads for the SMARCA4 KD or SMARCA4 heterozygous mutant datasets, respectively.

#### ATAC-seq data analysis

Both bulk ATAC-seq datasets were processed following the same method. Quality control was performed on raw data with FASTQC tool (v0.11.9). PEPATAC pipeline (v0.10.4) was executed to process raw reads and obtain accessible regions^46^. Briefly, raw reads were trimmed to remove adapters presence with Trimmomatic, trimmed reads were pre-aligned to mitochondrial genome and aligned to reference genome (hg38) with Bowtie2, deduplication was executed with Picard and accessible regions were called with Genrich discarding blacklisted regions (ENCODE hg38 v2). Pipeline default parameters were considered. All reference assembly resources were obtained from Refgenie asset manager^47^. Specifically, all required standard files were retrieved for hg38 alias (genome config file v0.4). Obtained .narrowPeak files from Genrich were imported into R (v4.4.3) to obtain a consensus peakset per genotype and condition (only for those clones with three replicates). Only regions with adjusted p-values (FDR) <0.05 and those overlapping in at least two samples (minimum 1bp in common) were considered accessible regions. For this purpose, findOverlapsOfPeaks function from ChIPpeakAnno R package (v3.40.0) was used with default parameters^48^. Peak annotation was performed with ChIPseeker R package (v1.42.1) considering a TSS region range between 5000bp upstream and 100bp downstream^49^. BigWig files were individually generated using deepTools (v3.5.1) bamCoverage with -binSize 10 -normalize CPM from the sorted deduplicated BAM files^50^. Differential accessibility analysis (DAA) was conducted with DESeq2 (v1.46.0)^51^. Accessible regions for testing were a union of all corresponding consensus peaksets by means of bedtools merge (maximum distance of 1bp, v2.30.0) of the concatenated and sorted list of regions^52^. Only canonical chromosomes were considered. To get a matrix of read counts per region, the function readCounts from csaw (v1.40.0) was used calling the deduplicated BAM files previously obtained^53^. For visualization purposes, counts were normalized by regularized logarithm method as implemented in DESeq2 R package (v1.46.0). Fitted statistical model included genotype (KD/WT or Mutant/WT depending on the dataset) and basal/treatment condition (only for the SMARCA4 mutant dataset). Additionally, an interaction between the two main effects was included in the model. Proper contrasts were defined to test the effect of genotype in both conditions (considering WT as reference) and the effect of treatment in both genotypes (considering basal condition as reference). In case of the SMARCA4 heterozygous mutant dataset, all four conditions were fitted simultaneously. Raw p-values were adjusted for multiple testing using the Benjamini-Hochberg False Discovery Rate (FDR). Differentially Accessible Regions (DARs) in any comparison were called with adjusted p-values (FDR) < 0.05 and absolute shrunken log2 Fold change > 1.

#### TF motif enrichment and footprinting analysis

DARs were interrogated for TF motif enrichment analysis which was conducted with STREME and TomTom tools from MEMESuite CLI (v5.5.2)^54^. First, STREME was executed for motif discovery with parameters --minw 8 -- maxw 20 --evalue --thresh 0.25 and considering DARs as primary sequence and the set of no differentially accessible regions as control sequence^55^. Secondly, discovered motifs were compared to HOCOMOCO v14 INVIVO (0-ranked based, ABC quality motif) database^56^ with TomTom and following parameters -min-overlap 5 -dist ed -thresh 0.25 -no-ssc^57^. Only significant results were considered for downstream interpretation (STREME E-value < 5e-02 and Tomtom q-value < 5e-02).

Same motif database was used to obtain TF deviation scores per sample by means of ChromVAR (v.1.28.0)^58^. Specifically, addGCBias, getBackgroundpeaks and computeDeviations functions were executed considering the same set of regions as in DAA (see ATAC-seq data analysis section) and hg38 reference genome. The matrix of deviation Z- scores was used for testing differential TF activity between conditions with limma (v.3.62.2)^59^. Fitted statistical model included one dummy variable by aggregating the genotype and basal/treatment condition. Motifs were ranked based on their logFC values and considered statistically significant with adjusted p-values < 0.01 (FDR).

TF footprinting analysis was conducted by means of TOBIAS (v0.14.0)^60^. Deduplicated BAM files of replicates from the same condition were merged into a unique alignment file. The same motif database as in the motif enrichment was used. ATACorrect, ScoreBigWig tools were executed sequentially with default parameters to obtain footprinting scores. Next, BINDetect tool was run to identify differentially bound motifs between conditions (pairwise comparisons). Differentially bound motifs in any comparison were called with p-values < 1e-100 and absolute differential TF footprinting change > 0.2.

#### Definition of AP-1-associated regulatory regions

AP-1-associated regions were defined as differentially accessible regions (DARs) containing a DNA motif of the AP-1 family motif cluster C_FOSL2_H14INVIVO.0.P.B (HOCOMOCO v14 INVIVO database; 0-ranked based, ABC quality motif), which groups the motifs for JUN, JUNB, JUND, ATF3, BATF2, FOS, FOSB, FOSL1 and FOSL2. Motif occurrences were those detected by TOBIAS (see TF footprinting analysis). Two subsets were defined according to the comparison used to call the DARs: AP- 1-associated regions with reduced basal accessibility in mutant cells (n = 1,373; downregulated DARs in T786I/WT versus WT under basal conditions), and MTX-inducible AP-1-associated regions remodeled in WT cells upon treatment (n = 750; upregulated DARs in MTX-treated versus untreated WT).

### RNA-SEQUENCING (RNA-seq)

#### Sample preparation

RNA-seq was performed on total RNA isolated from the same biological samples used for ATAC-seq. Three independent cultures of one SMARCA4 WT clone and one T786I/WT clone were collected under basal conditions or following treatment with 5 nM methotrexate (MTX) for 24 h. Total RNA was extracted from 1 × 10^6^ cells using the RNeasy Mini Kit (QIAGEN) according to the manufacturer’s instructions.

#### Library preparation and sequencing

RNA quality control, library preparation, and sequencing were performed at the CRG Genomics Unit (PRBB, Barcelona, Spain). RNA concentration and integrity were assessed using a Fragment Analyzer, and only samples with high RNA integrity (RQN ≥ 9.5) were used for poly(A) selection. In each experiment, all libraries were prepared simultaneously according to the facility’s standard protocol and total cDNA was sequenced using an Illumina NextSeq2000 sequencer (50 bp paired-end reads). Samples sequencing depth ranged between 52.5M and 63.7M reads.

#### RNA-seq data analysis

Quality control was performed on raw data with FASTQC tool (v0.11.9). Raw reads were trimmed to remove adapters presence with Trimgalore (v0.6.6)^61^. Default parameters were used except for a minimum quality of 15 (Phred score) and an adapter removal stringency of 3bp overlap. Trimmed reads were aligned to reference genome with STAR aligner tool (v2.7.8a)^62^. STAR was executed with default parameters except for the number of allowed mismatches which was set to 1. Required genome index was built with corresponding GRCh38 gtf and fasta files retrieved from Ensembl (http://ftp.ensembl.org/pub/release-106/). Obtained BAM files with uniquely mapped reads were considered for further analysis. Raw gene expression was quantified using featureCounts tool from subRead software (v2.0.1) with exon as feature^63^. Obtained raw counts matrix was imported into R Statistical Software environment (v4.4.3) for downstream analysis. Raw expression matrix included 61,552 genes. Bulk RNA-seq expression matrix included 12 samples: 3x SMARCA4 WT and 3x SMARCA4 heterozygous mutant per two conditions: basal and after 24 hours of methotrexate exposure. Prior to statistical analysis, those genes with less than 5 raw counts across the samples under test were removed. After pre-filtering, 26,728 genes were available for testing. For visualization purposes, counts were normalized by variance-stabilizing transformation method as implemented in DESeq2 R package (v1.46.0)^51^. Differential expression analysis (DEA) was conducted with DESeq2. Fitted statistical model included one dummy variable which was created aggregating the genotype and basal/treatment condition. Four different comparisons were tested: the effect of genotype in both conditions (considering WT as reference) and the effect of treatment in both genotypes (considering basal condition as reference). Obtained log2 fold change values were shrunken with apeglm shrinkage estimator R package (v1.28.0)^64^. Raw p-values were adjusted for multiple testing using the Benjamini-Hochberg False Discovery Rate (FDR). Differentially Expressed Genes (DEGs) in any comparison were called with adjusted p-values (FDR) < 0.05 and absolute shrunken log2 Fold change > 1.

#### Functional analysis

Pre-ranked Gene Set Enrichment Analysis (GSEA) was conducted over ranked list of genes based on the shrunken log2 Fold Change obtained from DEA. GSEA was conducted through fgseaMultilevel function from fgsea R package (v1.24.0) with default parameters (http://bioconductor.org/packages/fgsea/)^65^. GSEA interrogated MSigDB Hallmark database through msigdbr package (v7.5.1)^66^. Enrichment plots were generated with the same package. Overrepresented terms were called with adjusted p-value (FDR) < 0.1.

Overrepresentation analysis for selected sets of genes was conducted by means of clusterProfiler R package (v4.6.2)^67^ and interrogating CollecTRI regulon database^34^. Benjamini- Hochberg procedure was used to obtain adjusted p-values. Overrepresented TFs were called with adjusted p-value (FDR) < 0.1. The gene sets interrogated were: for Figure 3I, the leading- edge genes of the stress-response pathways enriched among genes downregulated in SMARCA4 T786I/WT versus WT cells under basal conditions (apoptosis, inflammatory response, interferon-α response, interferon-γ response, UV response DN, TGFβ response and epithelial-mesenchymal transition); and for Figure 5C, the stress-response and fitness signatures defined below.

#### Definition of stress-response and fitness signatures

Two custom gene signatures were derived from the GSEA of MTX-treated versus untreated WT cells (Figure 5B). The stress- response signature was built by pooling the leading-edge genes of the significantly enriched Hallmark pathways with positive NES that correspond to stress and damage-response programs (KRAS signaling, epithelial-mesenchymal transition, inflammatory response, IL2/STAT5 signaling, IL6/JAK/STAT3 signaling, TNFα signaling via NFκB, apoptosis and p53 pathway). The fitness signature was built by pooling the leading-edge genes of the significantly enriched pathways with negative NES that correspond to proliferative and biosynthetic programs (mitotic spindle, PI3K/AKT/mTOR signaling, unfolded protein response, mTORC1 signaling, E2F targets, oxidative phosphorylation, G2M checkpoint, MYC targets v1 and v2). The resulting signatures were then interrogated by pre-ranked GSEA on the comparison between MTX-treated SMARCA4 T786I/WT and MTX-treated WT cells (Figure 5D). The complete gene lists are provided in Supplementary Data 6.

### CHROMATIN IMMUNOPRECIPITATION (ChIP)-SEQUENCING AND ChIP-qPCR

#### Chromatin isolation and sonication

ChIP-seq was performed in three independent SMARCA4 WT clones and three T786I/WT clones cultured under basal conditions. For each sample, 10 million of cells were sequentially crosslinked with di-succynimidyl glutarate (DSG, Thermo-Fisher, 20593) and methanol-free formaldehyde (Pierce, 28906): 2 mM DSG for 1 h and formaldehyde for the final 15 min for SMARCA4 ChIP, as previously described^68^; 1 mM DSG for 15 min followed by 10 min formaldehyde for JUN ChIP. After crosslink quenching, nuclei were isolated by detergent-mediated lysis and resuspended in sonication buffer containing 1% SDS^69^. Chromatin was sheared in batches of 10 × 10⁶ cells to an average size of 200-500 bp by sonication (10 cycles of 30 s ON/30 s OFF at 4°C) using a Bioruptor Pico (Diagenode) and pre-cleared by adding 1% BSA, 12.5 µg/µl of salmon sperm DNA, 5 µg of rabbit IgG and protein A/G-Sepharose (GE Healthcare, 17-0618-01 and 17-0780-01). Pre- cleared chromatin was incubated for ∼16 h in RIPA buffer with 10 µg anti-SMARCA4 (Abcam ab110641, EPNCIR111A) per 20 × 10⁶ cell equivalents, 2 µg anti-JUN (Cell Signaling Technology, 9165) per 10 × 10⁶ cell equivalents, or the equivalent amount of irrelevant IgG, and precipitated with protein A/G-Sepharose. Crosslink was reversed overnight at 65°C and proteins were digested using Proteinase K (Roche) for 2 h at 55°C. DNA was purified using the MiniElute PCR Purification Kit (QIAGEN) and either submitted for library preparation and sequencing or used for quantitative PCR (ChIP-qPCR).

#### ChIP-qPCR

Fold enrichment was calculated relative to each sample input, normalized to irrelevant IgG, and background corrected by further normalization against an irrelevant negative control region (a gene desert in UCSC showing no enrichment in public JUN or SMARCA4 ChIP-seq or CUT&RUN datasets). Primers are listed in Appendix Table 2.

#### Library preparation and sequencing

DNA quality control, library preparation, and sequencing were performed at the CRG Genomics Unit (PRBB, Barcelona, Spain). Libraries from all samples were prepared simultaneously and sequenced on an Illumina NextSeq2000 platform (50 bp paired-end reads) according to the facility’s standard protocol. Samples sequencing depth ranged between 37.9M and 43.4M reads. In addition to the immunoprecipitated samples, one WT input and one T786I/WT input sample were generated by pooling equal amounts of input chromatin from the three corresponding biological replicates before library preparation.

#### ChIP-seq data analysis

Quality control was performed on raw data with FASTQC tool (v0.11.9). Raw reads were trimmed to remove adapters presence with Trimgalore (v0.6.6)^61^. Default parameters were used except for a minimum quality of 15 (Phred score) and an adapter removal stringency of 3bp overlap. Trimmed reads were aligned to reference genome with Bowtie2 (v2.4.4) which was executed parameters --no-mixed --end-to-end --no-discordant -- no-unal^70^. Required genome index was built with corresponding GRCh38 fasta file retrieved from Ensembl (release 106). Multimapped reads and those exhibiting MAPQ < 30 were removed. Duplicated reads were marked with SAMtools (v1.15)^71^. ENCODE blacklisted regions (hg38 v2) were removed prior to peak calling^72^. BigWig files were individually generated using deepTools (v3.5.1) bamCoverage with -binSize 10 -smoothLength 50 – effectiveGenomeSize 2701495711 -normalize RPGC^50^. The effective genome size was particularized to a read length of 50 bp and directly retrieved from deepTools web site (https://deeptools.readthedocs.io/en/develop/content/feature/effectiveGenomeSize.html). Peak calling was conducted by means of MACS2 (v2.2.7.1) with parameters -g 2701495711 - f BAMPE options^73^. The corresponding input sample was used in all peak calling computations. Peaks were called with adjusted p-values (FDR)<0.05. At this point, one of the three mutant samples was discarded for quality issues. Downstream analysis was conducted with R Statistical Software environment (v4.4.3). Peak annotation was performed with ChIPseeker R package (v1.42.1) considering a TSS region range between 5000bp upstream and 100bp downstream^49^. Required TxDb object was generated from the Ensembl GRCh38 release 106 gtf file retrieved. A consensus peakset was obtained for the SMARCA4 mutant condition. For this purpose, findOverlapsOfPeaks function from ChIPpeakAnno R package (v3.40.0) was used with default parameters^48^. For the SMARCA4 WT condition, peaks called in sample SW2 were used as representative of the condition, as this sample showed the highest peak recovery among WT replicates. The consensus peak set was defined as the union of the SW2 peak set and the SMARCA4 T786I/WT consensus peak set (n = 20,567 peaks). Tracks visualization were obtained by means of Integrative Genomics Browser (IGV) tool^74^.

Differential binding analysis (DBA) was conducted with DiffBind (v3.16.0)^75^. Default parameters were used except for the summits parameter which was set to 150bp in order to consider intervals of 300bp (between the minimum and first quartile peak width values). EdgeR was the statistical method used for all the three analyses. Differentially bound regions (DBRs) between SMARCA4 mutant and WT samples were called with adjusted p-values (FDR) < 0.05. Identified DBRs with any annotated gene were plotted with EnrichedHeatmap R package (v1.18.1). To summarize replicates per condition, normalizeToMatrix function was used based on corresponding BigWig files in w0 mean mode in 50bp windows. Tracks visualization were obtained by means of Integrative Genomics Browser (IGV)^74^.

#### Public CUT&RUN datasets

Publicly available RPMI8402 CUT&RUN datasets for histone modifications were obtained from the Gene Expression Omnibus (GEO) under accession GSE231963 (H3K27ac, GSM7306523; H3K4me1, GSM7306524; H3K4me3, GSM7306521).

## STATISTICAL ANALYSIS

GraphPad Prism 10 software and R software environment were used for statistical analysis. Statistical parameters, significance and the experimental unit (independent clone, independent experiment or individual animal) are indicated in the figures and legends. For in vitro proliferation, viability, drug-response, apoptosis and cell cycle assays, and qPCR analysis, statistical significance among groups was determined by Student’s t-test (data fitting normal distribution) or Mann–Whitney U test (data not fitting normal distribution) for two group comparison or one-way ANOVA with Tukey’s correction. For in vitro and in vivo competition assays, statistical significance was assessed using a two-tailed one-sample t-test against a theoretical value of 50%. Overlaps between sets of differentially accessible regions were evaluated using a hypergeometric test. Statistical thresholds applied to differential accessibility, differential expression, motif enrichment, footprinting and functional enrichment analyses are described in the corresponding Methods sections.

## CODE AVAILABILITY

This study did not involve the development of software. Scripts used to process the generated bulk RNA-seq, ChIP-seq and ATAC-seq data are available in Github repository: https://github.com/BigaSpinosaLab/PAPER_SMARCA4_Chemotherapy_Resistance_TALL.

## SEQUENCING DATA AVAILABILITY

Raw data from RNA-seq, ChIP-seq and ATAC-seq datasets generated in this study have been deposited in ArrayExpress/SRA repository under following accession numbers: E-MTAB-17364 (Bulk RNA-seq SMARCA4 heterozygous mutant and WT, basal and after methotrexate exposure conditions), E-MTAB-17367 (ChIP-seq SMARCA4 Mutant and WT, basal condition), E-MTAB-17401 (Bulk ATAC-seq SMARCA4 KD and WT, basal condition) and E-MTAB-17407 (Bulk ATAC-seq SMARCA4 heterozygous mutant and WT, basal and after methotrexate exposure conditions).

Supplementary data will be provided upon reasonable request.

**Appendix Table 1:**
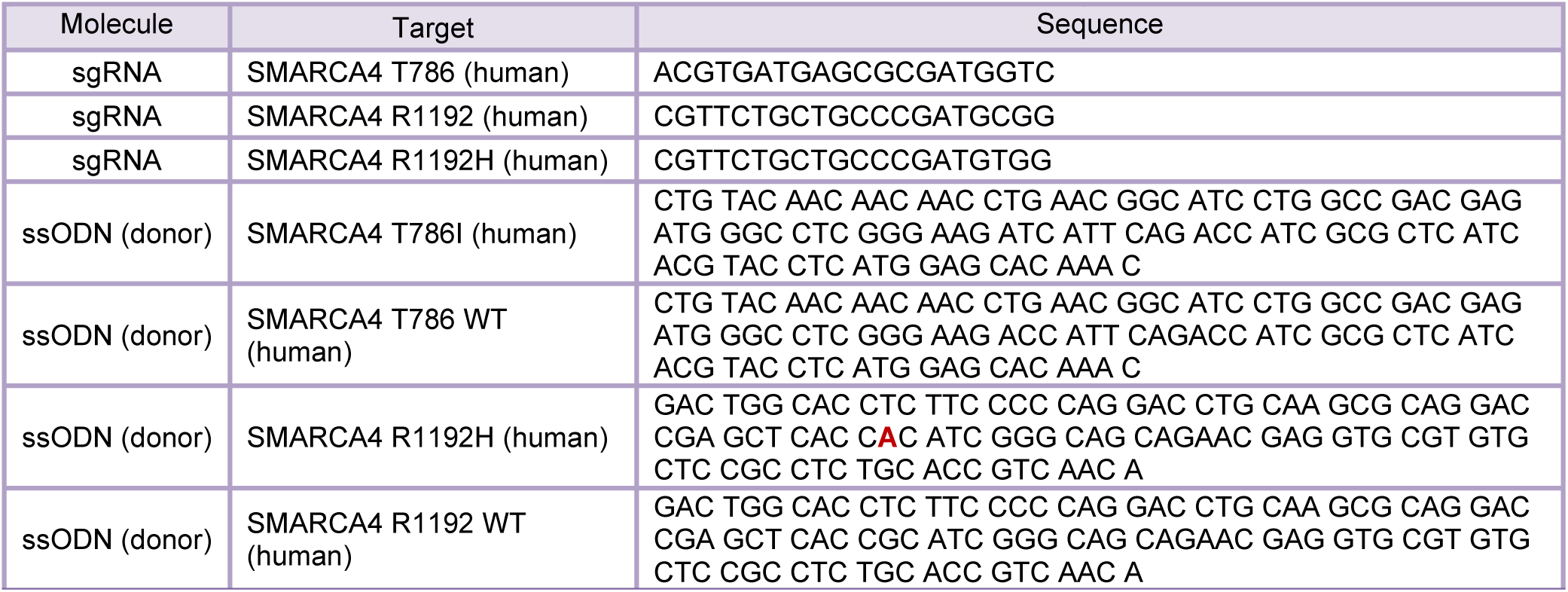
sgRNA and ssODN donor sequences.

**Appendix Table 2:**
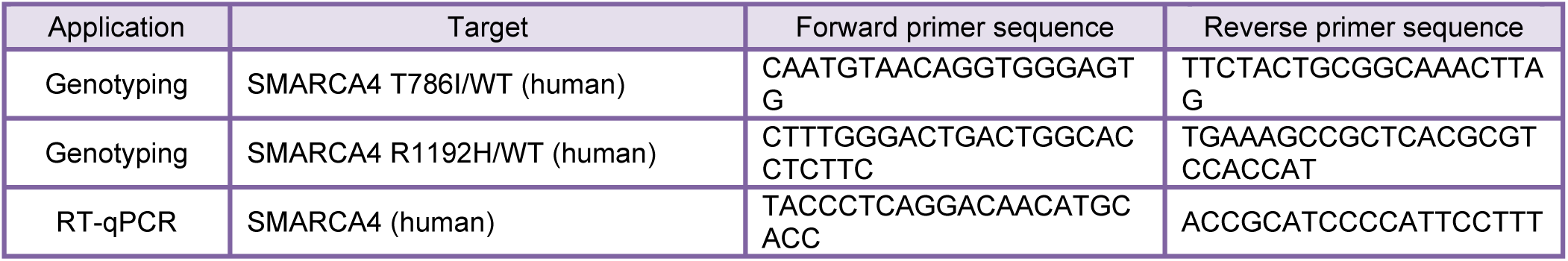
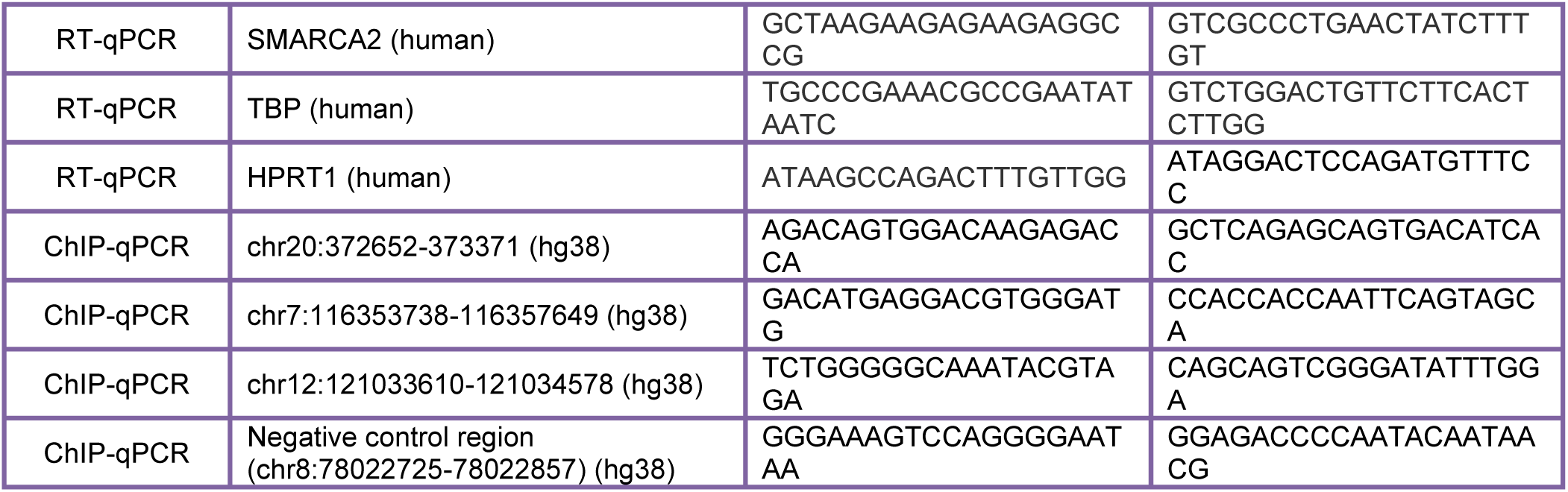
primer sequences.

**Appendix Table 3:** antibodies.

| Application | Target | Source | Reference (clone) | Host species | Dilution or µg |  |
|---|---|---|---|---|---|---|
| ChIP | SMARCA4 | Abcam | ab110641 (EPNCIR111A) | Rabbit | 10 µg / 20 million cells |  |
| ChIP | c-JUN | Cell Signaling | 9165 (60A8) | Rabbit | 2 µg / 10 million cells |  |
| WB | SMARCA4 | Cell Signaling | 49360S (D1Q7F) | Rabbit | 1/1,000 |  |
| WB | SMARCA2 | Cell Signaling | 48277 (D9E8B) | Rabbit | 1/1,000 |  |
| WB | α-Tubulin | Sigma | T6074 | Mouse | 1/10,000 |  |
| WB | Histone H3 | Abcam | ab1791 | Rabbit | 1/10,000 |  |
| Application | Target | Source | Reference (clone) | Target species | Fluorochrome | Dilution |
| Flow cytometry | CD45 | eBioscience | 45-0451-82 (30-F11) | Mouse | PerCP Cy5.5 | 1/200 |
| Flow cytometry | CD45 | BD Biosciences | 560178 (2D1) | Human | APC H7 | 1/200 |
| Flow cytometry | CD5 | Biolegend | 300622 (UCHT2) | Human | PE Cy7 | 1/200 |
| Flow cytometry | CD7 | BD Biosciences | 561604 (M-T701) | Human | APC | 1/200 |
| Flow cytometry | Ki67 | BD Biosciences | 558615 (B56) | Human | AF647 | 1/100 |

## Supporting information

merged supplemental figures and table

## ACKNOWLEDGEMENTS

The authors acknowledge all members of the Bigas, Espinosa and Bray laboratories for helpful discussions and technical support. We are grateful to Prof. Christoph Plass (DKFZ, Heidelberg) for critically reading the manuscript, to Prof. Alessandra Giorgetti and Dr. Damia Romero-Moya (Bellvitge Institute for Biomedical Research, IDIBELL, Barcelona) for providing support with initial genome editing designs, and to Prof. Luciano Di Croce (Center for Genomic Regulation, CRG, Barcelona) for kindly providing ATAC-seq reagents. We also thank the Center for Genomic Regulation (CRG) Genomics Unit, the Universitat Pompeu Fabra (UPF) Flow Cytometry Unit and the PRBB Animal Facility (MAPS) for their technical support. All illustrations in the manuscript were either original or created using BioRender.com. This work was supported by Agencia Estatal de Investigación (AEI) grant PID2022-137945OB-I00 and PID2025-168294OB-I00, José Carreras Leukäemie-Stiftung (DJCLS 14 R/2022 and DJCLS 21 R/2026), AGAUR and Generalitat de Catalunya (2021SGR39) and Fundación AECC (GC16173697BIGA) to AB, and by an MRC Programme Grant (MR/T014156/1) to SJB. Thanks to the contribution of Fundació Josep Carreras to AB. VGH and MM are recipients of a Sara Borrell fellowship (CD21/00145) and a research grant (grant number 377 CA22/00011), respectively, from the Instituto de Salud Carlos III (cofunded by the European Social Fund Plus and by the European Union). AV is a recipient of a Formación de Personal Investigador (FPI) PhD fellowship (PRE2022-000615) from the Spanish Ministry of Science, Innovation and Universities.

## AUTHOR CONTRIBUTION

AB, LE, and VGH conceptualized the study and experimental design; VGH, GF, AV, JG, AI, MAM and EP performed experiments; MM designed, performed and supervised bioinformatic analysis; CB performed bioinformatic analysis; FB and SJB provided resources and supervision. AB and VGH analyzed data, supervised the research and wrote the manuscript with the input from all authors; AB provided resources.

