## Supplementary material for "SMARCA4 MUTATIONS PROMOTE CHEMOTHERAPY RESISTANCE IN T-ACUTE LYMPHOBLASTIC LEUKEMIA THROUGH ALTERED CHROMATIN REMODELING AT AP-1-ASSOCIATED REGULATORY REGIONS": merged supplemental figures and table

**Figure S1. Generation and characterization of isogenic SMARCA4-mutant T-ALL models.**

**A.** Western blot analysis of SMARCA4 and SMARCA2 protein expression in cytoplasmic and nuclear fractions across T-ALL cell lines. Tubulin and H3 detection were used as loading and fractionation controls.

**B.** Representative Sanger sequencing chromatograms confirming introduction of heterozygous T786I and R1192H mutations in RPMI8402 cells and reversion of the endogenous R1192H mutation in Jurkat cells.

**C.** Western blot analysis of SMARCA4 and SMARCA2 protein expression in nuclear extracts from RPMI8402 and Jurkat WT and edited clones.

**D.** Cell proliferation of WT and SMARCA4-mutant RPMI8402 clones under basal culture conditions.

**E.** Cell proliferation upon SMARCA4 knockdown or control shRNA (shControl) in RPMI8402 cells using two independent shRNAs targeting SMARCA4. Asterisks indicate the comparison of each SMARCA4-targeting shRNA with shControl at the corresponding time point.

**F.** Cell viability of WT and SMARCA4-mutant RPMI8402 clones under basal conditions.

**G.** Cell viability upon SMARCA4 knockdown in RPMI8402 cells. Asterisks indicate the comparison of each SMARCA4-targeting shRNA with shControl at the corresponding time point.

**H.** Cell cycle analysis of WT and SMARCA4 T786I/WT RPMI8402 clones under basal conditions. Left, quantification of cell-cycle phase distribution determined by DAPI and Ki67 staining in four independent clones per genotype. Each dot represents an independent clone, with values calculated as the mean of two independent experiments. Right, representative flow cytometry density plots.

**I.** Sensitivity of RPMI8402 WT and T786I/WT clones to etoposide and doxorubicin. Graphs represent the area under the curve (AUC) quantification of independent isogenic clones.

**J.** Western blot analysis of nuclear SMARCA4 protein levels following SMARCA4 knockdown in RPMI8402 cells using two independent shRNAs targeting SMARCA4.

**K**. Chemotherapy response of shControl and SMARCA4-knockdown RPMI8402 cells following treatment with MTX, VDL, or etoposide.

For panels D-G, each curve corresponds to an independent clone and data represent mean ± SD from 3 (RPMI8402) or 2 (Jurkat) independent experiments. Statistical significance was determined by one-way ANOVA with Tukey's correction for multiple comparison testing.

For panel H, each dot represents an independent clone, with values calculated as the mean of two independent experiments. Statistical significance between WT and SMARCA4-mutant groups was assessed using two-tailed unpaired t-tests for each cell-cycle phase.

For panel I, each dot represents an independent isogenic clone. Values for each clone were calculated as the mean of three independent experiments. Statistical significance was assessed using two-tailed unpaired t-tests.

****$p < 0.0001$, ***$p < 0.001$, **$p < 0.01$, *$p < 0.05$.

**Figure S2. Additional validation of the competitive advantage conferred by relapse-associated SMARCA4 mutations.**

**A.** In vitro competition assays using three independent GFP-labeled SMARCA4 T786I/WT RPMI8402 clones, each competed against an equicellular pool of unlabeled WT clones, during MTX (top) or VDL (bottom) treatment.

**B.** Reciprocal labeling strategy, in which a GFP-labeled pool of WT clones was competed against an unlabeled pool of SMARCA4 T786I/WT clones, during MTX (left) or VDL (right) treatment.

**C.** Relative contribution of WT and SMARCA4 T786I/WT RPMI8402 cells within the human leukemic compartment in the spleen and PB of chemotherapy-treated and untreated mice. The dashed line indicates the equal representation of both populations (50%) as at the start of the experiment.

**D.** Experimental design of the in vitro and in vivo competition assays performed using Jurkat SMARCA4 R1192H/WT (unedited) and reverted WT clones.

**E.** Relative abundance of Jurkat SMARCA4 R1192H/WT (unedited) and reverted WT populations during MTX treatment, VDL treatment, or untreated culture. Data represents the mean ± SD of 3 independent experiments.

**F.** Percentage of Jurkat cells in bone marrow (BM), liver, spleen and peripheral blood (PB) at sacrifice. Jurkat cells identified as mouse CD45- human CD45+CD5+CD7+ by flow cytometry and gated on DAPI negative cells. Each dot represents an independent animal.

**G.** Relative contribution of WT and SMARCA4 R1192H/WT Jurkat cells within the human leukemic compartment of the indicated tissues in chemotherapy-treated and untreated mice. The dashed line indicates the equal representation of both populations (50%) as at the start of the experiment.

For panels A-B, data represent the mean ± SD of three independent experiments per mix.

For panel E, data represent the mean ± SD of 3 independent experiments per mix.

For panels F-G, n = 8 animals per group; 4 per labeling strategy. Only animals with leukemic engraftment above the predefined cut-off were included in panel G.

Statistical significance was assessed using a two-tailed one-sample t-test against a theoretical value of 50%. ****p < 0.0001, ***p < 0.001, **p < 0.01, *p < 0.05.

**Figure S3. SMARCA4 hypomorphic alterations selectively impair AP-1-associated chromatin accessibility.**

**A.** Volcano plot showing differential chromatin accessibility following SMARCA4 knockdown in RPMI8402 cells determined by ATAC-seq. Differentially accessible regions (DARs) were defined as absolute shrunken log2FC > 1 and adjusted p-value < 0.05. Total regions analyzed, n = 61,037; DARs with reduced accessibility, n = 9,308; DARs with increased accessibility, n = 1,384.

**B–C.** Venn diagrams showing the overlap between DARs identified in SMARCA4 knockdown RPMI8402 cells (vs shControl) and DARs identified in SMARCA4 T786I/WT RPMI8402 cells (vs WT). Statistical significance was calculated using a hypergeometric test.

**D.** Genomic annotation of regions showing reduced or increased accessibility following SMARCA4 knockdown in RPMI8402 cells, as defined in A.

**E.** SMARCA4 ChIP-seq occupancy profiles at the SMARCA4 consensus peakset (union of WT and mutant peaks, n = 20,567) in three independent WT and two independent SMARCA4 T786I/WT RPMI8402 clones. Signals are shown within a ±2 kb window centered on each peak.

**F.** Transcription factor motif enrichment analysis of regions displaying reduced (top) or increased (bottom) accessibility following SMARCA4 knockdown in RPMI8402 cells. ATAC-seq regions without significant accessibility changes were used as the background set. Statistical significance thresholds were STREME E-value < $5 e^{-02}$ and Tomtom q-value < $5 e^{-02}$ . For each motif, the indicated number and percentage correspond to DARs containing the respective motif

**G.** Comparison of differential transcription factor occupancy inferred by footprinting analysis in SMARCA4 T786I/WT cells (versus WT) and in SMARCA4-knockdown cells (versus shControl). Transcription factors are colored according to whether the change in occupancy was significant in both conditions, only in mutant cells or only in knockdown cells. Statistical significance threshold: absolute ΔTF occupancy > 0.2 and p-value < 1 e-100. AP-1 family members are shown in bold.

Statistical thresholds used for differential accessibility and expression analyses are indicated in Methods.

**Figure S4. Global chromatin responses to methotrexate treatment.**

**A.** Venn diagram showing the overlap between regions gaining accessibility after MTX treatment in WT and SMARCA4 T786I/WT cells. Regions losing accessibility are not represented, as only 8 were identified in SMARCA4 T786I/WT cells.

**B.** Principal component analysis (PCA) of ATAC-seq samples before and after MTX treatment. The plot includes three biological replicates from one WT and one SMARCA4 T786I/WT clone used for differential accessibility analyses, together with one additional WT and one SMARCA4 T786I/WT clone used for validation.

**C**. Transcription factor motif enrichment analysis of regions gaining accessibility following MTX treatment. ATAC-seq regions without significant accessibility changes were used as the background set. Statistical significance thresholds were STREME E-value < $5 e^{-02}$ and Tomtom q-value < $5 e^{-02}$. For each motif, the indicated number and percentage correspond to DARs containing the respective motif.

Statistical thresholds used for differential accessibility analyses are described in Methods.

**Figure S5. Expression of AP-1 family members and effect of pharmacological AP-1 inhibition in RPMI8402 cells.**

**A.** VST-normalized expression of AP-1 family members in WT and SMARCA4 T786I/WT RPMI8402 cells under basal conditions and after MTX treatment (24 h, 5 nM). Each dot represents an independent replicate. Data from RNA-seq experiments in WT and mutant clone #1. Asterisks indicate the adjusted p-value from the differential expression analysis for the indicated comparison (DESeq2; see Methods).

**B.** Area under the curve (AUC) quantification corresponding to the dose-response curves of parental RPMI8402 cells represented in Figure 6C. Data are shown as mean ± SD of 3 independent experiments. Statistical significance of AUC values between conditions was assessed using two-tailed unpaired t-tests.

**C.** Dose-response curve of RPMI8402 cells treated for 72 h with the AP-1 inhibitor T-5224. Data represent mean ± SD.

Statistical thresholds used for differential expression analyses are indicated in Methods. ****p < 0.0001, ***p < 0.001, **p < 0.01, *p < 0.05.

### Table S1

#### SMARCA4 mutations detected in relapsed T-ALL cohorts

<table>
<tr><th>Cohort</th><th>Age group</th><th>Patient ID</th><th>SMARCA4 mutation</th><th>VAF Dx (%)</th><th>VAF Rel (%)</th><th>Stage</th></tr>
<tr><td rowspan="4">Sentís et al.<br>(n = 19)<br><br>González-Gil et al.<br>(n = 37)</td><td rowspan="4">Adult</td><td>PAT8</td><td>T786I</td><td>ND</td><td>34.62</td><td>Rel</td></tr>
<tr><td>PAT14</td><td>G911V</td><td>ND</td><td>8.72</td><td>Rel</td></tr>
<tr><td>PAT14</td><td>T912S</td><td>ND</td><td>10.00</td><td>Rel</td></tr>
<tr><td>PAT14</td><td>G1162S</td><td>ND</td><td>40.31</td><td>Rel</td></tr>
<tr><td rowspan="4">González-Gil et al.<br>(n = 37)</td><td rowspan="4">Adult</td><td>PAT25</td><td>R1243Q</td><td>ND</td><td>4.52</td><td>Rel</td></tr>
<tr><td>PAT32</td><td>H1191R</td><td>ND</td><td>61.15</td><td>Rel</td></tr>
<tr><td>PAT36</td><td>R425Q</td><td>ND</td><td>43.96</td><td>Rel</td></tr>
<tr><td>PAT36</td><td>D1506W</td><td>ND</td><td>2.56</td><td>Rel</td></tr>
<tr><td rowspan="2">Oshima et al.<br>(n = 31)</td><td rowspan="2">Pediatric</td><td>Pat27</td><td>M1040R</td><td>na</td><td>na</td><td>Shared</td></tr>
<tr><td>Pat27</td><td>R1243W</td><td>na</td><td>na</td><td>Dx</td></tr>
<tr><td rowspan="3">Li et al.<br>(n = 16)</td><td rowspan="3">Pediatric</td><td>SJTALL 013801</td><td>E882D</td><td>ND</td><td>32.14</td><td>Rel</td></tr>
<tr><td>SJALL 043861</td><td>P974H</td><td>ND</td><td>18.18</td><td>Rel</td></tr>
<tr><td>SJALL 043857</td><td>R1192H</td><td>ND</td><td>16.33</td><td>Rel</td></tr>
<tr><td>Kunz et al.<br>(n = 13)</td><td>Pediatric</td><td>S00207</td><td>R966W</td><td>0.83</td><td>58.36</td><td>Rel</td></tr>
</table>

The 19 adult cases from Sentís et al. are included within the 37 cases of González-Gil et al., giving a non-redundant cohort of n = 97. The overlapping cases were analysed by whole-genome sequencing in Sentís et al. and by a NGS panel in González-Gil et al.; mutations detected by either approach are reported here.

### FIGURE S1

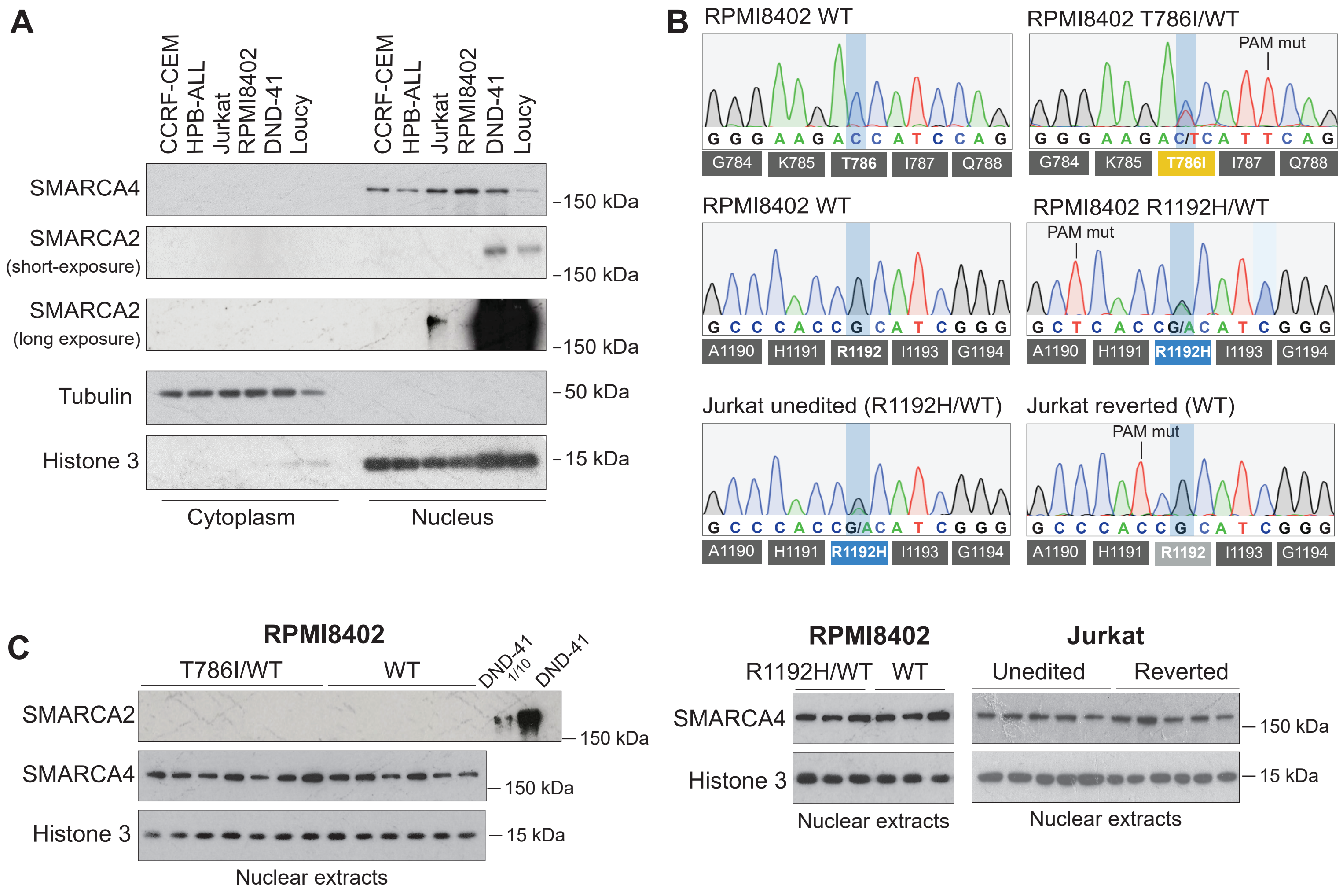

A
CCRF-CEM
HPB-ALL
Jurkat
RPMI8402
DND-41
Loucy
SMARCA4
SMARCA2
(short-exposure)
SMARCA2
(long exposure)
Tubulin
Histone 3
150 kDa
50 kDa
15 kDa
Cytoplasm
Nucleus
B
RPMI8402 WT
RPMI8402 T786I/WT
PAM mut
G784
K785
T786
T786I
I787
Q788
RPMI8402 R1192H/WT
A1190
H1191
R1192
R1192H
I1193
G1194
Jurkat unedited (R1192H/WT)
Jurkat reverted (WT)
C
RPMI8402
T786I/WT
WT
DND-41 1/10
DND-41
Nuclear extracts
R1192H/WT
Jurkat
Unedited
Reverted

### FIGURE S1

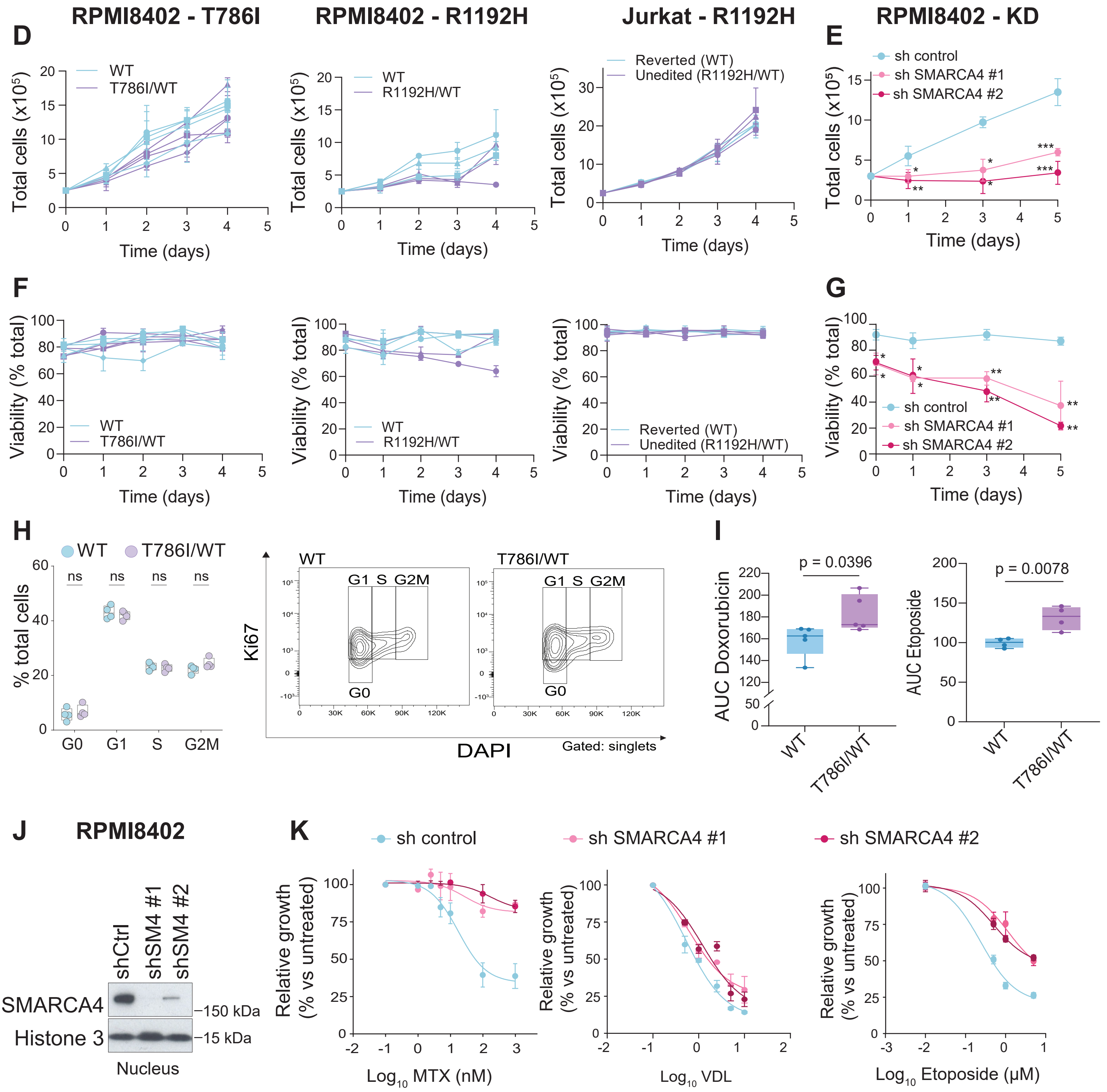

RPMI8402 - T786I
RPMI8402 - R1192H
Jurkat - R1192H
RPMI8402 - KD
D
E
F
G
H
I
J
K
Total cells (x10^5)
Time (days)
WT
T786I/WT
R1192H/WT
Reverted (WT)
Unedited (R1192H/WT)
sh control
sh SMARCA4 #1
sh SMARCA4 #2
Viability (% total)
% total cells
G0
G1
S
G2M
ns
Ki67
DAPI
Gated: singlets
AUC Doxorubicin
AUC Etoposide
p = 0.0396
p = 0.0078
RPMI8402
shCtrl
shSM4 #1
shSM4 #2
SMARCA4
Histone 3
150 kDa
15 kDa
Nucleus
Relative growth (% vs untreated)
Log10 MTX (nM)
Log10 VDL
Log10 Etoposide (μM)

### FIGURE S2

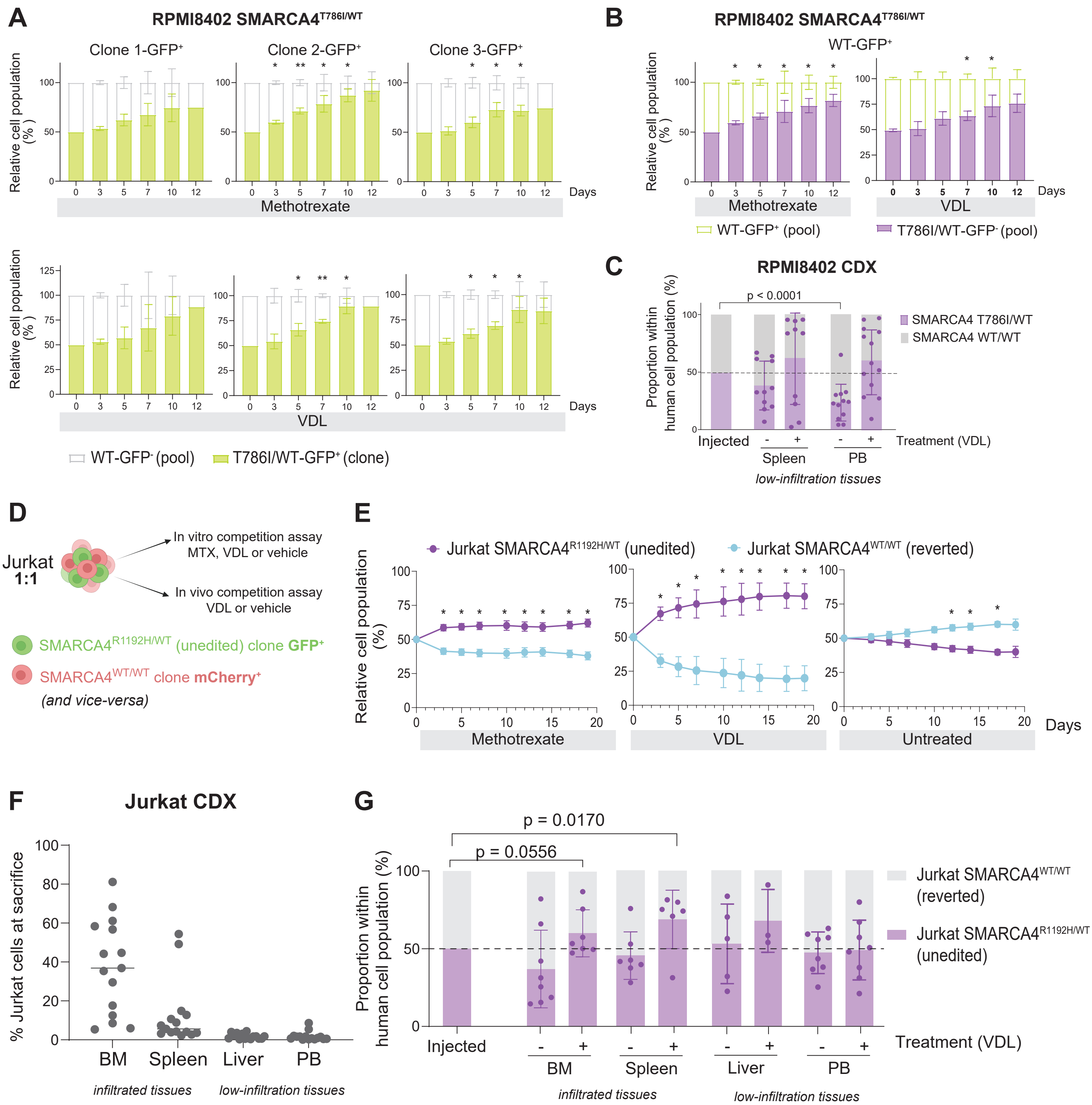

A
RPMI8402 SMARCA4T786I/WT
Clone 1-GFP+
Clone 2-GFP+
Clone 3-GFP+
Relative cell population (%)
Methotrexate
VDL
Days
WT-GFP- (pool)
T786I/WT-GFP+ (clone)
B
RPMI8402 SMARCA4T786I/WT
WT-GFP+
Methotrexate
VDL
WT-GFP+ (pool)
T786I/WT-GFP- (pool)
C
RPMI8402 CDX
p < 0.0001
Proportion within human cell population (%)
SMARCA4 T786I/WT
SMARCA4 WT/WT
Injected
Spleen
PB
Treatment (VDL)
low-infiltration tissues
D
Jurkat 1:1
In vitro competition assay MTX, VDL or vehicle
In vivo competition assay VDL or vehicle
SMARCA4R1192H/WT (unedited) clone GFP+
SMARCA4WT/WT clone mCherry+
(and vice-versa)
E
Jurkat SMARCA4R1192H/WT (unedited)
Jurkat SMARCA4WT/WT (reverted)
Relative cell population (%)
Methotrexate
VDL
Untreated
Days
F
Jurkat CDX
% Jurkat cells at sacrifice
BM
Spleen
Liver
PB
infiltrated tissues
low-infiltration tissues
G
p = 0.0170
p = 0.0556
Proportion within human cell population (%)
Jurkat SMARCA4WT/WT (reverted)
Jurkat SMARCA4R1192H/WT (unedited)
Injected
BM
Spleen
Liver
PB
Treatment (VDL)
infiltrated tissues
low-infiltration tissues

### FIGURE S3

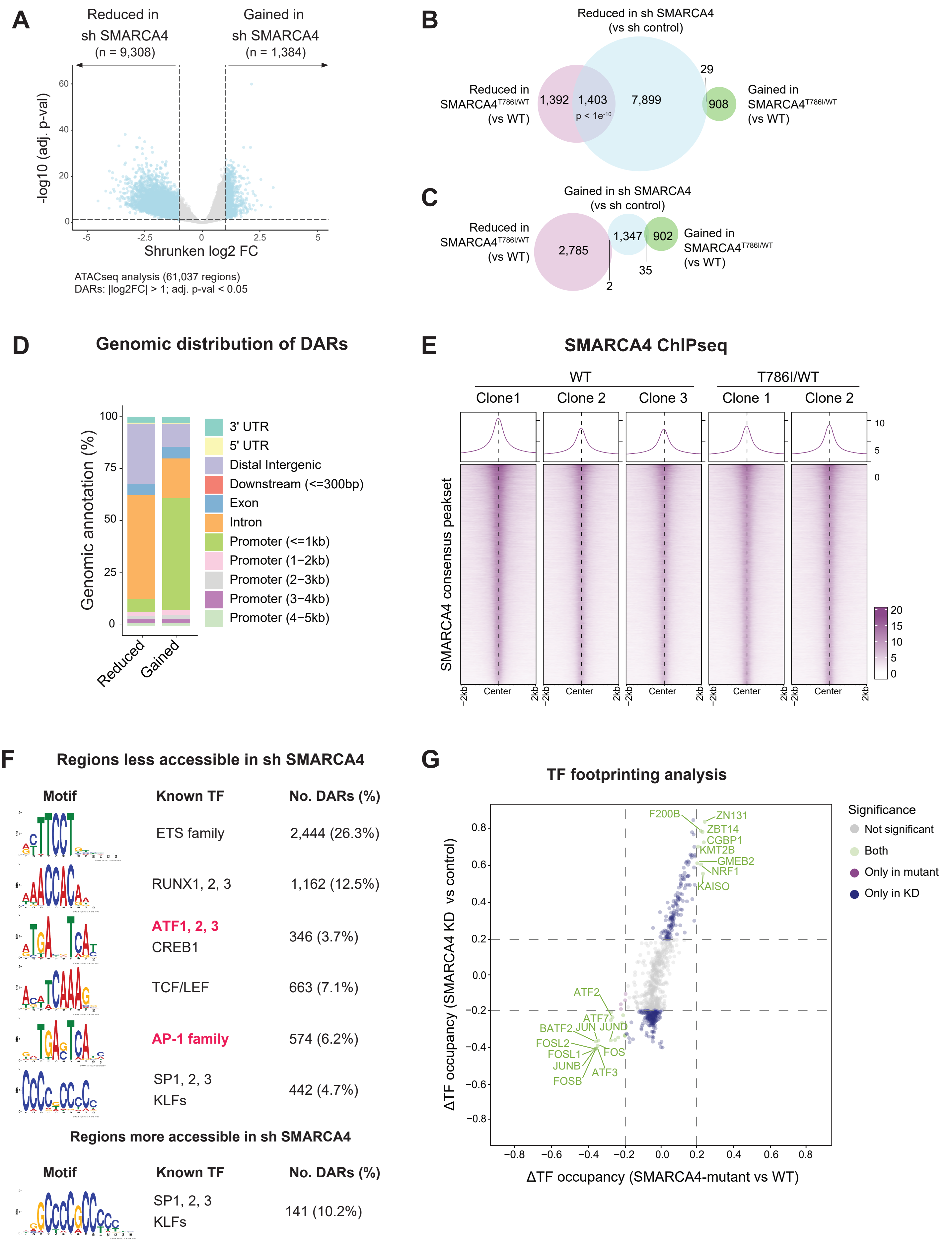
A
Reduced in sh SMARCA4 (n = 9,308)
Gained in sh SMARCA4 (n = 1,384)
-log10 (adj. p-val)
Shrunken log2 FC
ATACseq analysis (61,037 regions)
DARs: |log2FC| > 1; adj. p-val < 0.05
B
Reduced in sh SMARCA4 (vs sh control)
Reduced in SMARCA4T786I/WT (vs WT)
Gained in SMARCA4T786I/WT (vs WT)
1,392
1,403
p < 1e-10
7,899
29
908
C
Gained in sh SMARCA4 (vs sh control)
Reduced in SMARCA4T786I/WT (vs WT)
Gained in SMARCA4T786I/WT (vs WT)
2,785
1,347
902
35
2
D
Genomic distribution of DARs
Genomic annotation (%)
Reduced
Gained
3' UTR
5' UTR
Distal Intergenic
Downstream (<=300bp)
Exon
Intron
Promoter (<=1kb)
Promoter (1−2kb)
Promoter (2−3kb)
Promoter (3−4kb)
Promoter (4−5kb)
E
SMARCA4 ChIPseq
WT
T786I/WT
Clone1
Clone 2
Clone 3
Clone 1
Clone 2
SMARCA4 consensus peakset
-2kb
Center
2kb
F
Regions less accessible in sh SMARCA4
Motif
Known TF
No. DARs (%)
ETS family
2,444 (26.3%)
RUNX1, 2, 3
1,162 (12.5%)
ATF1, 2, 3
CREB1
346 (3.7%)
TCF/LEF
663 (7.1%)
AP-1 family
574 (6.2%)
SP1, 2, 3
KLFs
442 (4.7%)
Regions more accessible in sh SMARCA4
Motif
Known TF
No. DARs (%)
SP1, 2, 3
KLFs
141 (10.2%)
G
TF footprinting analysis
ΔTF occupancy (SMARCA4 KD vs control)
ΔTF occupancy (SMARCA4-mutant vs WT)
Significance
Not significant
Both
Only in mutant
Only in KD
F200B
ZN131
ZBT14
CGBP1
KMT2B
GMEB2
NRF1
KAISO
ATF2
ATF7
BATF2
JUN
JUND
FOSL2
FOSL1
FOS
JUNB
ATF3
FOSB

### FIGURE S4

**A** More accessible sites following MTX short-treatment

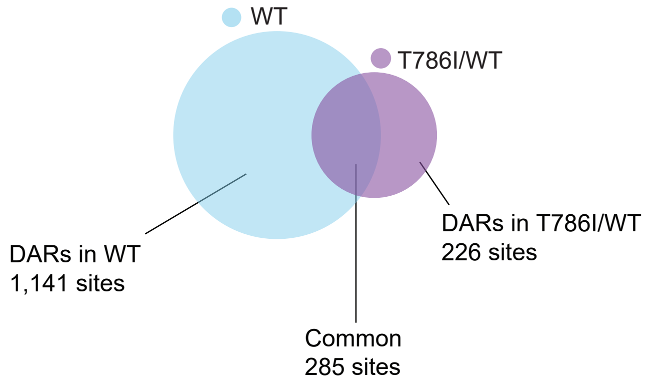


**B**

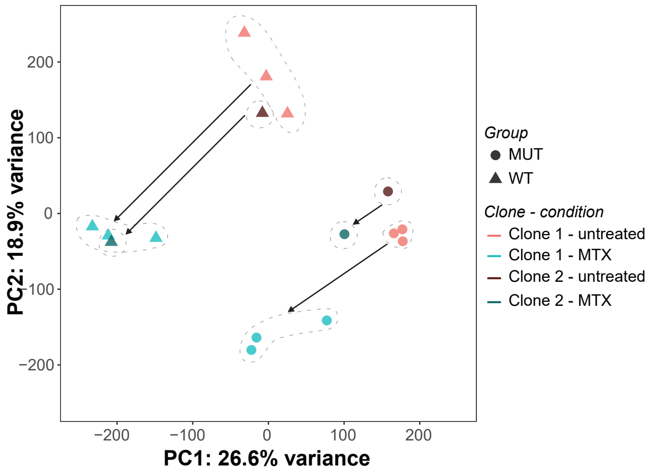


**C** **Regions gaining accessiblity after 24h-MTX**

| **Motif** | **Known TF** | **No. DARs (%)** |
|---|---|---|
| 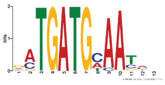 | **AP-1 family**<br>CEBPA,B,G<br>CREB5,NFIL3 | 324 (19.6%) |
| 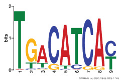 | **AP-1 family**<br>CREB5 | 579(35.5%) |

Other enriched motifs: ZN560, ZN121, ZN441

### FIGURE S5

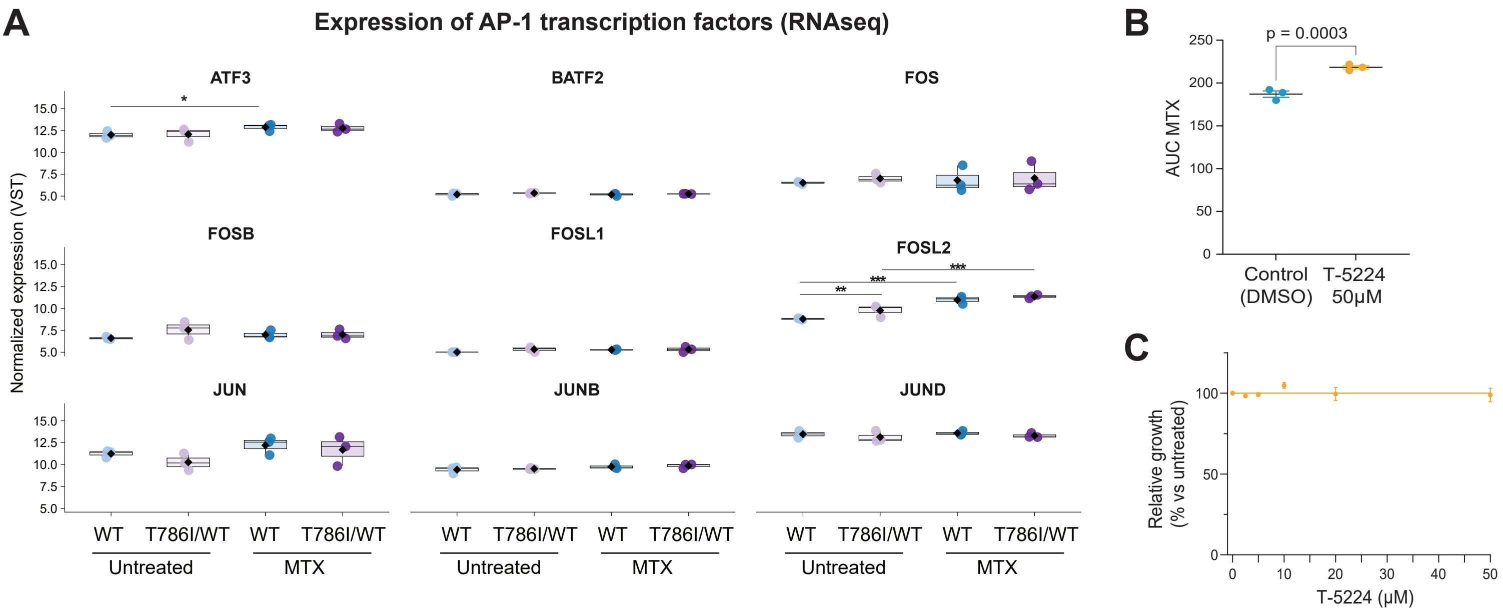
A
Expression of AP-1 transcription factors (RNAseq)
ATF3
BATF2
FOS
FOSB
FOSL1
FOSL2
JUN
JUNB
JUND
Normalized expression (VST)
15.0
12.5
10.0
7.5
5.0
*
**
***
WT
T786I/WT
Untreated
MTX
B
p = 0.0003
AUC MTX
250
200
150
100
50
0
Control (DMSO)
T-5224 50μM
C
Relative growth (% vs untreated)
100
50
0
10
20
30
40
50
T-5224 (μM)
